# Upregulation of breathing rate during running exercise by central locomotor circuits

**DOI:** 10.1101/2021.07.28.453983

**Authors:** Coralie Hérent, Séverine Diem, Gilles Fortin, Julien Bouvier

## Abstract

While respiratory adaptation to exercise is compulsory to cope with the increased metabolic supply to body tissues and with the necessary clearing of metabolic waste, the neural apparatus at stake remains poorly identified. Using viral tracing, *ex vivo* and *in vivo* optogenetic and chemogenetic interference strategies in mice, we unravel interactive locomotor and respiratory networks’ nodes that mediate the respiratory rate increase that accompanies a running exercise. We show that the mesencephalic locomotor region (MLR) and the lumbar spinal locomotor pattern generator (lumbar CPG), which respectively initiate and execute the locomotor behavior, access the respiratory network through distinct entry points. The MLR directly projects onto the inspiratory rhythm generator, the preBötzinger complex (preBötC), and can trigger a moderate increase of respiratory frequency, prior to, or even in the absence of, locomotion. In contrast, the lumbar CPG projects onto the retrotrapezoid nucleus (RTN) that in turn contacts the preBötC to enforce, during effective locomotion, higher respiratory frequencies. These data expand, on the one hand, the functional implications of the MLR beyond locomotor initiation to a *bona fide* respiratory modulation. On the other hand, they expand the adaptive respiratory ambitions of the RTN beyond chemoception to “locomotor- ception”.

## Introduction

Breathing is a vital behavior which must combine extreme robustness with continuous adaptability. One striking example is the abrupt augmentation of ventilation at the transition from rest to running in order to maintain homeostasis in spite of increased metabolic demand (DiMarco et al., 1983; Duffin and Bechbache, 1983; Mateika and Duffin, 1995). This “exercise hyperpnoea” is manifested by an increase in both respiratory frequency and volume. It has long been proposed that its main trigger, at least for acute exercise, is of neuronal nature, i.e. relies on activatory signals from locomotor effectors or circuits impacting onto the respiratory generator in the brainstem (Mateika and Duffin, 1995; Gariepy et al., 2010; Duffin, 2014; Paterson, 2014). However, the underlying cells and circuits are not fully elucidated.

We recently uncovered that running hyperpnoea can, at least in mice but very likely in all quadrupeds, occur without temporal synchronization of breathes to strides (Hérent et al., 2020). This, together with the ventilatory response seen in men during mental simulation of exercise (Tobin et al., 1986; Decety et al., 1991; Decety et al., 1993) or in the absence of peripheral signals (Fernandes et al., 1990), highlights that the main neuronal trigger of running hyperpnoea is of central, rather than peripheral, origin. Of particular interest are therefore brain regions that command or execute locomotor movements and could provide a parallel drive to respiratory centers (Eldridge et al., 1981; Eldridge et al., 1985). The mesencephalic locomotor region (MLR) in the dorsal midbrain is considered the main site of locomotor initiation throughout the animal kingdom (Shik et al., 1966; Le Ray et al., 2011; Ryczko et al., 2016; Chang et al., 2021) likely including humans (Jahn et al., 2008; Sébille et al., 2019). Stimulation of the MLR, and particularly its cuneiform nucleus (CnF) component, engages forward locomotion at a speed that is commensurate to the intensity of the stimulus (Bachmann et al., 2013; Roseberry et al., 2016; Caggiano et al., 2018; Josset et al., 2018; Dautan et al., 2020; van der Zouwen et al., 2021), making it a candidate neuronal encoder and driver of running intensity. The possibility that the MLR may provide a parallel activation of respiratory centers is suggested by work in the lamprey (Gariepy et al., 2011), an ancestral vertebrate specie, but this has yet not been investigated in terrestrial mammals. Another central drive to respiratory centers may originate in the circuits of the spinal cord that elaborate the locomotor rhythm and coordinate the motor output during ongoing locomotor movements, often referred to as a “Central Pattern Generator” or CPG (Grillner, 2006; Kiehn, 2016; Grillner and El Manira, 2020). Indeed, pharmacological activation of the lumbar enlargement, where the hindlimb CPG circuit is thought to reside, can upregulate the frequency of respiratory-like activities on *ex vivo* preparations from neonatal rats (Le Gal et al., 2014; Le Gal et al., 2020). While this is suggestive of ascending projections to respiratory centers, the underlying circuit and its functionality during running has not been documented.

Another gap of knowledge resides in the identification of the respiratory neurons targeted by descending (e.g., MLR) or ascending (e.g., from the CPG) locomotor drives. In mammals, the respiratory rhythm is paced by a confined cluster of neurons in the ventromedial medulla, the pre-Bötzinger complex (preBötC, Smith et al., 1991; Del Negro et al., 2018). Direct activation or inactivation of preBötC glutamatergic neurons respectively increases, or reduces and even arrests, respiratory rate (Tan et al., 2008; Alsahafi et al., 2015; Cui et al., 2016; Vann et al., 2018). The preBötC receives inputs from several brain areas including the midbrain (Yang et al., 2020) and, in the lamprey, MLR neurons were shown to contact a presumed homologue of the preBötC (Mutolo et al., 2010; Gariepy et al., 2012). This makes the preBötC a candidate for promptly entraining respiration during exercise in mammals. More rostrally, an area collectively referred to as the parafacial (pF) respiratory region may be another candidate of respiratory regulation during metabolic challenges including effort. In particular in this region, non-catecholaminergic *Phox2b*-expressing neurons (defining the Retrotrapezoid Nucleus, RTN) are well-known for their capacity to rapidly upregulate respiratory rate in the context of central CO2 chemoception (Abbott et al., 2009; Abbott et al., 2011). They might also support active expiration which is thought to accompany exercise (Ainsworth et al., 1989; Iscoe, 1998; Abraham et al., 2002). *Phox2b-*positive neurons in the pF region have been shown to be activated during running (Barna et al., 2012, 2014), during locomotor-like activity on *ex vivo* neonatal rat preparations (Le Gal et al., 2014), and their chemogenetic silencing limits exercise capacity in running rats (Huckstepp et al., 2015).

Here we sought to investigate the central circuits interfacing locomotor and respiratory centers in the resourceful mouse model. We found the existence of both a descending drive from the MLR, and of an ascending drive from the locomotor CPG of the lumbar spinal cord. Remarkably, the MLR is capable of upregulating breathing rate even before the initiation of actual limb movements. We further uncovered that the two systems both have access to respiratory rhythm generation mechanisms albeit through two different synaptic schemes. The MLR directly projects to the preBötC, but not to the pF respiratory region, while the lumbar spinal cord targets the RTN which in turns contacts the preBötC. Our work therefore demonstrates two locomotor central drives that may underlie breathing adaptability during running and their synaptic nodes in the respiratory central network.

## Results

### Glutamatergic CnF neurons project to the preBötC

We first examined whether locomotor-promoting MLR neurons in mice, as is the case with the lamprey (Gariepy et al., 2012), contact neuronal groups involved in respiratory rhythm generation. The MLR contains two major subdivisions, the cuneiform nucleus (CnF) containing glutamatergic (Glut+) neurons, and the pedunculopontine nucleus (PPN) containing both glutamatergic and cholinergic neurons (Roseberry et al., 2016; Caggiano et al., 2018; Josset et al., 2018). Since locomotor initiation is mostly attributed to the former, we traced the projections of CnF neurons by unilateral stereotaxic injections of a Cre-dependent Adeno Associated Virus (AAV) coding the fluorescent protein eYFP in *Vglut2^Cre^* adult mice (Vong et al., 2011) (Figure 1a, b). Abundant eYFP-positive fibers were detected in the preBötC, defined as located ventrally to the nucleus ambiguus, containing SST-positive neurons and between antero-posterior levels -7.0 and -7.4 from bregma (Stornetta et al., 2003; Tan et al., 2008). These projections were found bilaterally with an ipsilateral predominance (Figure 1c, d, g). In contrast, projections were very sparse in the pF respiratory area, defined as located immediately ventral, ventro-median and ventro-lateral to the facial motor nucleus (7N, between antero-posterior levels -6.5 and -5.7 from Bregma, Figure 1e-g). We also observed CnF projections to the parabrachial nucleus and to the nucleus of the tractus solitarius (data not shown). To verify that CnF neurons synaptically target preBötC neurons, we made use of a genetically restricted two-virus approach (Kim et al., 2016) to reveal preBötC putative inputs anatomically. Functional preBötC neurons can be efficiently delineated as Glut+ neurons with commissural projections (Koshiya and Smith, 1999; Bouvier et al., 2010). We therefore drove the expression of the TVA receptor and the rabies protein G using a retrogradely transported Cre-dependent Herpes Simplex Virus (HSV, Neve et al., 2005; Reinhard et al., 2019) injected in the preBötC on one side. Seven days later, a G-deleted and EnvA-pseudotyped rabies (Rb) virus coding mCherry was injected in the preBötC on the other side (Figure 1h). As demonstrated previously, this leads to the expression of the Rb virus in projection-defined neuronal somata (Usseglio et al., 2020), here commissural Glut+ neurons of the preBötC (Figure 1i), and, from these starter cells, to its transynaptic spread to upstream neurons. Transynaptically labelled neurons, i.e., expressing only the Rb-driven fluorophore, were detected in the CnF and PPN nuclei bilaterally (Figure 1j). Putative inputs to Glut+ preBötC neurons were also detected in the contralateral preBötC, the periaqueductal grey, the superior colliculus and the NTS, and only few cells were detected in the pF respiratory region on either side (data not shown, Figure S1). Altogether, these converging anterograde and retrograde tracings demonstrate that Glut+ CnF neurons make direct contact with candidate respiratory rhythm generating neurons in the preBötC, but not in the pF respiratory region.

**Figure 1:**
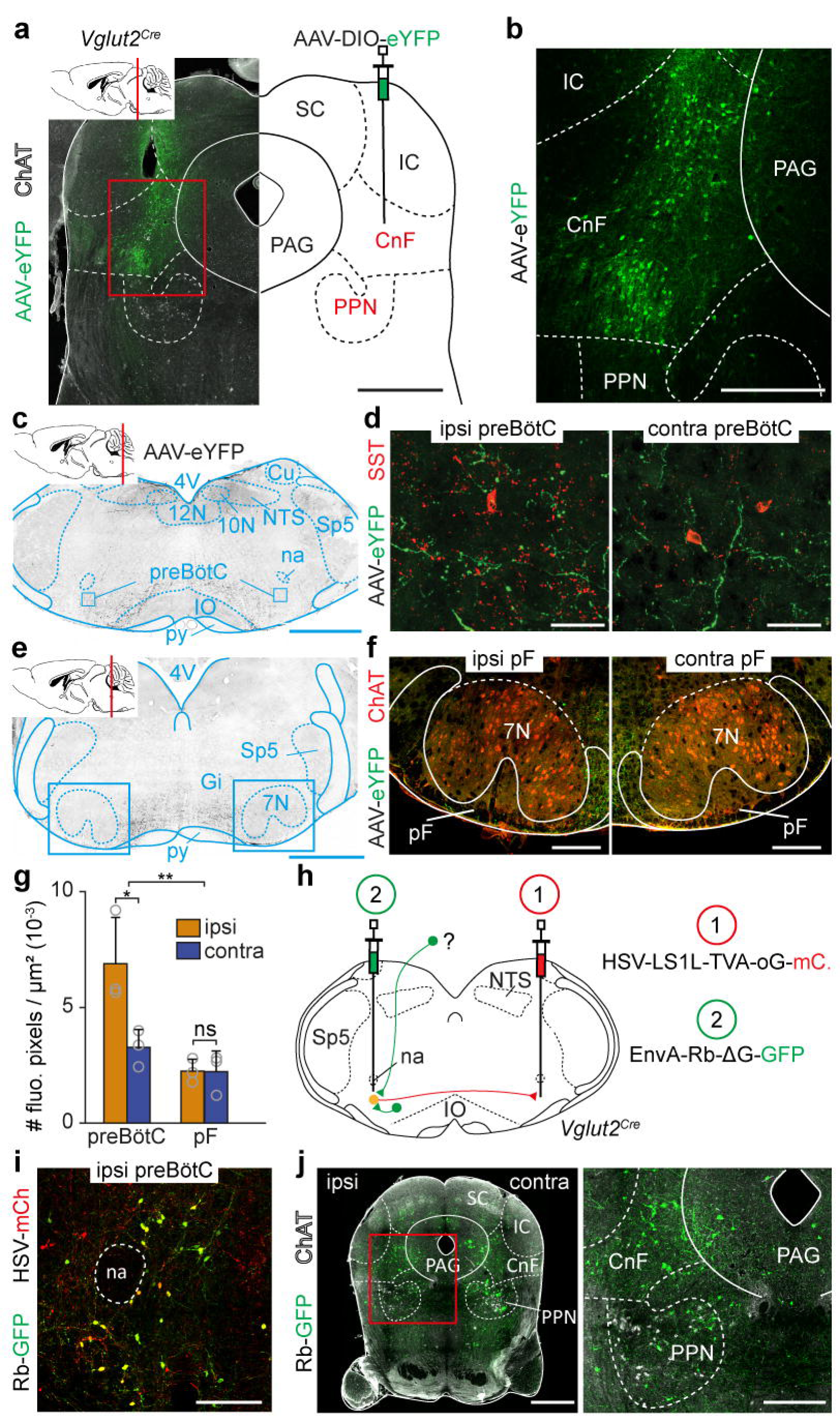
CnF glutamatergic neurons contact the preBötC inspiratory generator. **(a)** Transverse section at the level of the injection site in the mesencephalon, showing glutamatergic CnF neurons transfected unilaterally with an AAV-DIO-eYFP on a *Vglut2^Cre^* adult mouse. The MLR component nuclei CnF and PPN are in red. Cholinergic PPN neurons are identified by Choline Acetyl Transferase (ChAT) expression. Scale bar, 1 mm. **(b)** Magnification of the boxed area in (**a**), showing transfected eYFP^+^ cells concentrated in the CnF. Scale bar, 400 µm. **(c)** Transverse section in the caudal medulla showing the distribution of CnF projections in the ventral reticular formation including the preBötC. Scale bar, 1 mm. **(d)** Magnifications of the boxed areas in (**c**), over the ipsilateral and contralateral preBötC containing SST^+^ cells and processes. Note that the CnF projects to both sides with an ipsilateral predominance. Scale bar, 40 µm. **(e)** Transverse section in the rostral medulla showing the distribution of CnF projections at the level of the parafacial (pF) respiratory area. Scale bar, 1 mm. **(f)** Magnifications of the red boxed areas over the ipsilateral and contralateral pF respiratory areas, delineated as ventral, ventro-median and ventrolateral to the ChAT-positive facial motor nucleus (7N). Note that the CnF projects very little to the pF area on either side. Scale bar, 250 µm. Panels (**a-f**) are representative of n=3 animals. **(g)** Bar-graphs showing the mean density ± SD across mice of eYFP^+^ fluorescent pixels located ipsilaterally and contralaterally in the preBötC and the pF. Grey open circles are the mean values of individual mice. **, p < 0.01; *, p < 0.05; ns, not-significant (Wilcoxon matched-pairs tests; preBötC, 9 sections per side; pF, 9 sections per side; from n = 3 mice). **(h)** Retrograde transynaptic monosynaptic tracing from glutamatergic commissural preBötC neurons. An HSV-LS1L-TVA- oG-mCherry is first injected in the contralateral preBötC followed by a G-deleted and EnvA pseudotyped Rb virus (EnvA- ΔG-Rb-GFP) in the ipsilateral preBötC in a *Vglut2^Cre^* adult mouse. **(i)** Magnification over the ipsilateral preBötC from a transverse section showing double-transfected “starter” cells. Scale bar, 200 µm. **(j)** Left: Transverse section at the MLR level showing putative presynaptic cells (Rb-GFP^+^) on a ChAT background. Scale bar, 1 mm. Right: magnification of the boxed area showing the presence of transynaptically labelled cells in the CnF and PPN. Scale bar, 250 µm. Panels (**h,j**) are representative of n= 3 animals. See also Figure S1. Abbreviations used in all figures: PAG: periaqueductal gray; IC: inferior colliculus; SC: superior colliculus; PPN: pedunculopontine nucleus; CnF: cuneiform nucleus; 4V: fourth ventricle; 10N: dorsal motor nucleus of vagus ; 12N: hypoglossal motor nucleus; NTS: nucleus tractus solitarius; py: pyramidal tract; IO: inferior olive; Cu: cuneate nucleus; na: nucleus ambiguus; preBötC: pre-Bötzinger complex; pF: parafacial respiratory area; Sp5: spinal trigeminal nucleus; 7N: facial nucleus; Gi: gigantocellular reticular nucleus.

### Glutamatergic CnF neurons modulate inspiratory rhythm generation mechanisms

We next tested by optogenetics the ability of Glut+ CNF neurons to functionally impact the preBötC. One hallmark of preBötC responses to phasic incoming inputs is their phase-dependency to the ongoing rhythm (Cui et al., 2016; Baertsch et al., 2018). To evaluate this for CnF-evoked preBötC responses, we virally delivered the light-activated excitatory opsin Channelrhodopsin 2 (ChR2) in the CnF on one side (AAV-DIO- ChR2-eYFP) of *Vglut2^Cre^* adult mice, and implanted an optic fiber over the injection site (Figure 2a). Breathing cycles were measured in awake animals with whole body plethysmography (WBP, (DeLorme and Moss, 2002), Figure 2b) while short single-pulse (50 ms) photoactivations were delivered randomly during the respiratory cycle. We calculated the resultant phase shift, expressed as the perturbed cycle duration over the control cycle duration as done previously (Cui et al., 2016; Baertsch et al., 2018, Figure 2b). A phase shift <1 (perturbed cycle duration lower than the control one) indicates a shortening of the respiratory cycle, a phase shift >1 (perturbed cycle duration higher than the control one) indicates a lengthening, and a phase shift equal to 1 (perturbed cycle duration equal to the control one) indicates no effect. We found that unilateral photostimulation of CnF neurons elicited an ectopic inspiratory burst and shortened the respiratory cycle (Figure 2c) but that this effect was dependent on the timing of light-activation during the respiratory cycle. Specifically, the most significant cycle shortening was seen when delivering light-pulses during early expiration (phase: 0.5 - 0.6, phase shift: 0.80 ± 0.10, p < 0.0001 Figure 2c). In contrast, the effect was only minimal when photostimulations were delivered in late expiration. Intriguingly, in addition to the immediate inspiratory response to photostimulation, we observed a shortening (although less drastic) of the next two consecutive cycles suggesting that activation of the CnF also determines a longer lasting modulatory action on preBötC rhythm generation (Figure S2a-d). No alteration of the respiratory cycle was however seen in mock trials in control mice that do not express ChR2 (Figure S3a, b), ruling out a contribution of light-evoked temperature changes locally (Stujenske et al., 2015). To ascertain that this modulation of respiratory rhythm generation owes to direct projections of Glut+ CnF neurons to the preBötC, we next aimed at photo- activating ChR2-expressing fibers in the preBötC following the delivery of the ChR2 virus in the CnF (Figure 2d). This led to a similar phase-dependent shortening of the respiratory cycle, with again a maximal shortening observed when light-activations are delivered in early expiration (phase: 0.5 - 0.6; phase shift: 0.79 ± 0.15, p < 0.0001, Figure 2e, f). This effect disappeared after the 2^nd^ subsequent respiratory cycle (θ+3, Figure S2e-h). Light deliveries in the preBötC of control mice that do not express ChR2 did not produce any noticeable effect (Figure S3d, e). Overall, this indicates that the direct projections of the Glut+ CnF neurons indeed conform to phase-dependent activation of preBötC neurons, highlighting excitatory modulations of inspiratory burst generation.

**Figure 2:**
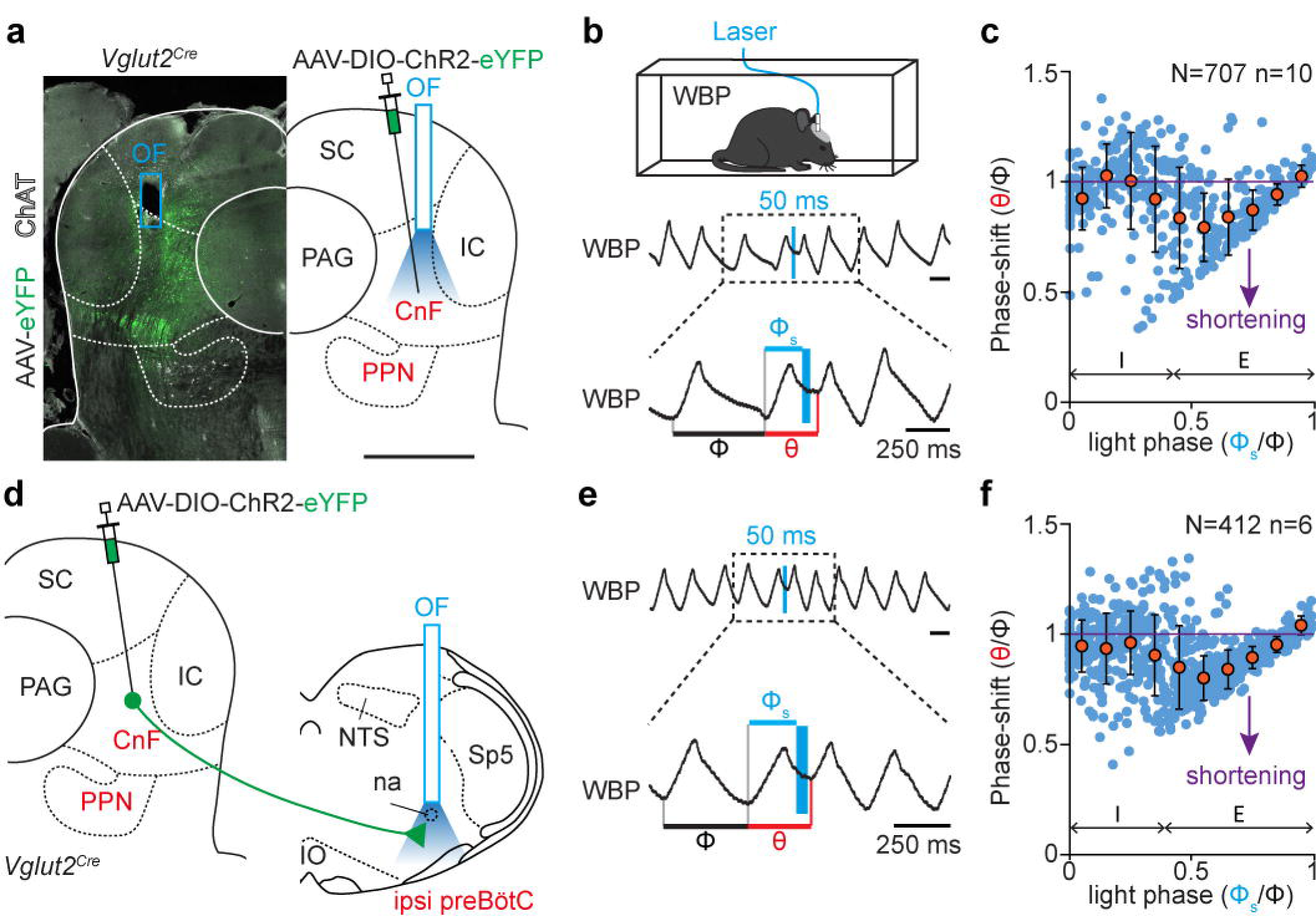
Photoactivation of glutamatergic CnF neurons impacts respiratory rhythm generation in the preBötC. **(a)** Left: low magnification of a transverse section at the level of the injection and optic fiber (OF) implantation sites, showing glutamatergic CnF neurons transfected with an AAV-DIO-ChR2-eYFP in a *Vglut2^Cre^* adult mouse. The MLR component nuclei CnF and PPN are in red. Cholinergic PPN neurons are identified by Choline Acetyl Transferase (ChAT) expression. Scale bar, 1 mm. **(b)** Top: setup for recording ventilation using whole-body plethysmography (WBP) during optogenetic activations. Middle: WBP recordings around a single 50 ms light-stimulation pulse of glutamatergic CnF neurons during the expiratory phase of one respiratory cycle (inspirations are upwards, expirations are downwards). Note that the stimulation shortens the respiratory cycle. Bottom: magnification of the above trace showing the control cycle (φ, black), the phase of light-simulation (φs, blue), and the perturbed cycle (θ, red). **(c)** Plot of the phase-shift (perturbed cycle normalized to the control cycle: θ/φ) as a function of the phase of light- stimulation normalized to the control cycle (φs/φ). Values < 1 (purple line) indicate a shortening of the perturbed cycle. Note that the phase-shift is shortened when the light pulse occurs in the expiratory phase of the respiratory cycle. Inspiration (I) and expiration (E) mean durations are indicated. Blue circles represent individual data from N = 707 random trials from n = 10 mice. Orange circles are averages ± SD across all trials within 0.1 ms bins. **(d)** Experimental strategy for photostimulating glutamatergic CnF neurons fibers in the preBötC: the injection is done as above but a second optic fiber is implanted above the preBötC on the ipsilateral side. **(e-f)** Same as in (**c-d**) during a single 50 ms light-stimulation of the preBötC area showing again a shortening of the respiratory cycle when the light pulse is delivered during expiration. N = 412 random trials from n = 6 mice. See also Figures S2, S3.

### Glutamatergic CnF neurons modulate breathing in synergy with locomotion

Through their access to rhythm generating neurons in the preBötC, Glut+ CnF neurons might be capable of upregulating breathing frequency in synergy with locomotor initiation. To access respiratory parameters during vigorous displacement movements, we made use of our recently-developed method for chronic electromyographic (EMG) recordings of the diaphragm, the main inspiratory muscle (Hérent et al., 2020). Animals were thus made to express ChR2 in Glut+ CnF neurons as above, EMG-implanted, and placed in a linear corridor. Light was delivered in trains of 1 s duration at increasing pulse frequencies when animals were stationary at one end of the corridor (Figure 3). Animals were filmed from the side and their displacement speed computed using markerless video-tracking (Mathis et al., 2018) as performed previously (Hérent et al., 2020). In line with numerous studies (Roseberry et al., 2016; Caggiano et al., 2018; Josset et al., 2018; Dautan et al., 2020), we found that photoactivation of Glut+ CnF neurons at 15 Hz or more engages animals in forward locomotion (Figure 3a), and that higher stimulation frequencies impose faster regimes, shorten the delay between light onset and locomotor initiation and increase the occurrence of left-right synchronous gaits (Figure S4). Importantly, CnF photostimulations were associated with an increased respiratory rate (Figure 3b, c), an effect that was not seen in control mice that do not express ChR2 (Figure S3c). In what we consider a remarkable observation, during CnF photoactivations that effectively engage running, respiratory rate increased in a two-step sequential manner. In a first step, that we term the “pre- loco” phase, a modest increase was seen immediately at light onset but before the first locomotor movements (i.e., during the delay between light onset and the initiation of locomotion, Figure 3d). The mean respiratory rate during this “pre-loco” phase was found significantly higher than baseline but was not correlated to the photostimulation frequency (Figure 3e). In a second step, when the animals effectively engage in locomotion (“loco” phase), the respiratory rate was further augmented (Figure 3d). There, respiratory rate was still not strongly dependent on stimulation frequency, however it was proportional to the actual displacement speed (Figure 3f). This likely reflects the variability of locomotor velocities at a given stimulation frequency (Figure S4b). However, we found that during CnF photoactivations, respiratory and locomotor rhythms were not temporally synchronized (Figure S5), in line with what we recently reported during spontaneous running in mice (Hérent et al., 2020). These results indicate that respiratory frequency during CnF-evoked locomotion is upregulated immediately at light onset and before the initiation of locomotion, and further upregulated during actual locomotion yet without a phasic relation to limb cycles.

**Figure 3:**
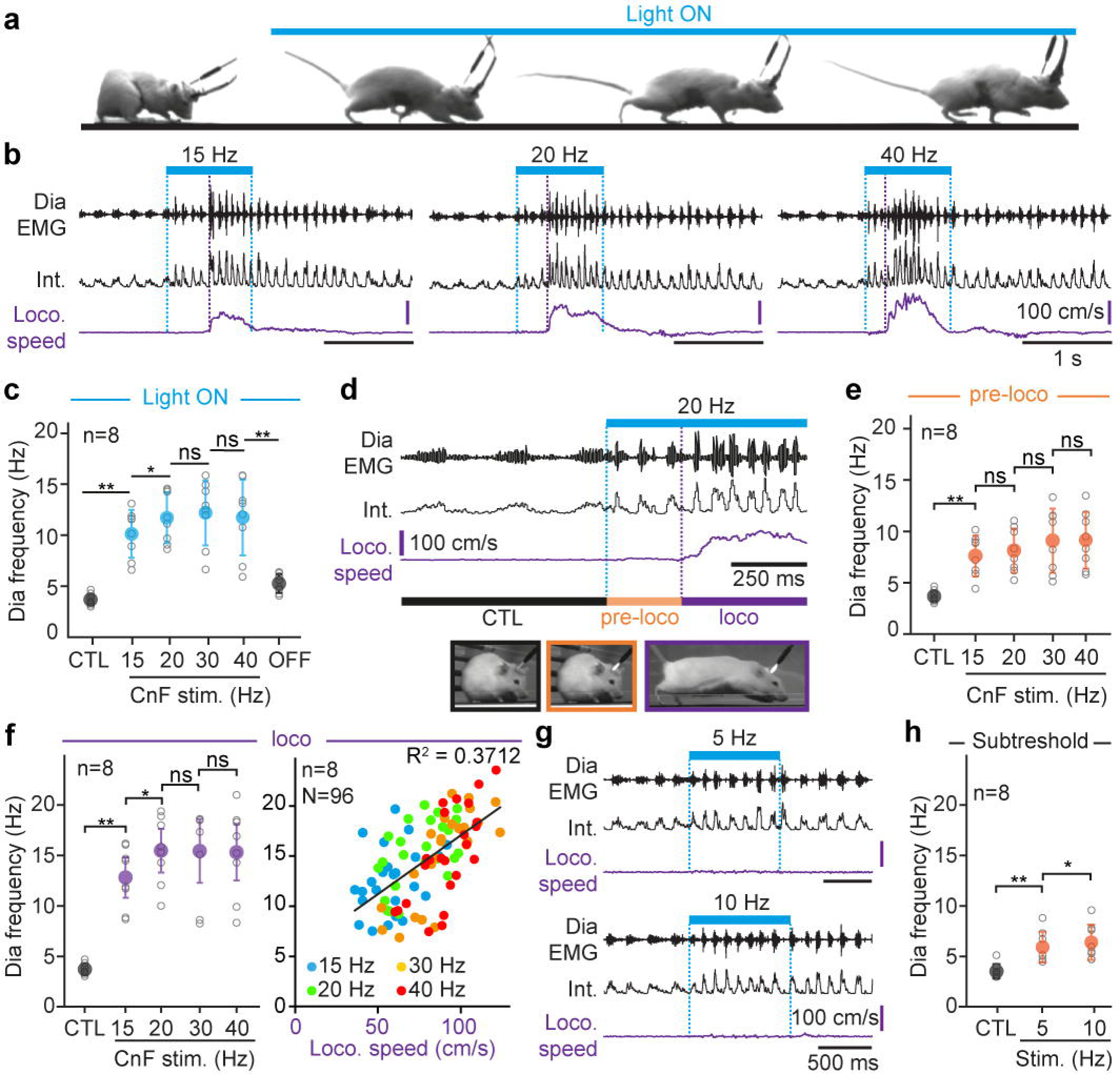
Glutamatergic CnF neurons upregulate breathing in synergy with, and even in the absence of, locomotion. **(a)** Example snapshots showing that photostimulation of the glutamatergic CnF neurons for 1 s (20 Hz pulses) triggers running. The injection and implantation strategy is similar to that described in Figure 2a. **(b)** Raw (Dia EMG) and integrated (Int.) electromyogram of the diaphragm and instantaneous locomotor speed during 1 s CnF photostimulations at 15, 20 and 40 Hz. Dotted blue lines: onset and offset of light stimulations; purple dotted line: onset of the locomotor episode. Note that breathing frequency increases at light onset, before the animal starts running, and increases further upon running. **(c)** Quantification of the diaphragmatic frequency at rest (CTL), during the entire CnF photostimulation at increasing frequencies, and following light offset (OFF). **(d)** Top: enlarged EMG recording of one representative animal during rest (CTL) and during the 1 s CnF photostimulation at 20 Hz which is divided in two phases: “pre-loco” (orange) during which the animal remains still, and “loco” (purple) when the animal engages in running. Bottom: snapshots of one animal in the 3 states. **(e)** Quantification of the diaphragm frequency in control (CTL) and during the “pre-loco” phase at increasing CnF photostimulation frequencies showing a significant increase from baseline for 15 Hz photostimulations but no further increase at higher frequencies. **(f)** Left: similar quantifications during the “loco” phase. Right: color plot showing the diaphragm frequency in relation to locomotor speed. Note that 37 % of the diaphragm frequency correlates with the locomotor speed (linear regression, black line). N = 96 trials from n = 8 mice. **(g)** Example traces from one representative animal during CnF photostimulations at 5 and 10 Hz. Although these stimulations are below the threshold for running initiation, breathing frequency is significantly increased. **(h)** Quantifications of the diaphragm frequency in control (CTL) and during subthreshold CnF stimulations. In all graphs, gray open circles are the means of 3 trials for individual animals and colored circles are the means ± SD across n mice. ns, not significant. *, p < 0.05, **, p < 0.01 (Wilcoxon matched-pairs tests). See also Figures S3, S4 and S5.

The former observation above suggests that Glut+ CnF neurons can modulate respiratory activity independently of their action on limb movements, in line with what was reported in the lamprey (Gariepy et al., 2012). To demonstrate this further, we reduced the intensity of photostimulations below the threshold for locomotor initiation (5 and 10 Hz, Figure 3g, Figure S4b). We found that, in spite of absent locomotor movements, the respiratory frequency was indeed significantly increased from baseline (Figure 3h). Altogether, these analyses demonstrate that i) Glut+ CnF neurons can upregulate breathing before, or even in the absence of, locomotor movements, ii) during CnF-evoked locomotion, the highest increase in breathing rate from rest occurs when actual locomotor movements are engaged, iii) respiratory frequency increase during the “loco” phase are proportional to the displacement speed, and iv) breaths are not phase-locked to cyclic limb movements.

### The spinal locomotor circuits project to the pF respiratory region

From the above observation we reasoned that the engagement in actual locomotor movements may be associated with a stronger activating drive onto respiratory centers which could originate in executive lumbar locomotor circuits. We thus examined the ascending projections to the brainstem of glutamatergic lumbar neurons that are essential in the regulation of locomotor speed (Kjaerulff and Kiehn, 1996; Hagglund et al., 2010; Hagglund et al., 2013; Talpalar et al., 2013). To do so, we injected a Cre-dependent AAV-eYFP vector bilaterally in the ventral laminae of the 2^nd^ lumbar segment (L2) of adult *Vglut2^Cre^* animals (Figure 4a) and examined projections in the brainstem reticular formation. In contrast to the anterograde tracings from the CnF, this revealed very few, if any, eYFP-positive fibers in the preBötC but their abundant presence in the pF respiratory region (Figure 4b-f). To discriminate passing fibers from putative synaptic contacts, we first repeated these spinal cord injections with an AAV that drives a presynaptic, synaptophysin-fused, GFP (AAV-DIO-Syp-GFP) and indeed observed GFP-positive puncta in the pF (Figure 4g). We also performed similar spinal injections this time with a high-titer AAV coding the Cre recombinase (AAV1-Syn-Cre, Zingg et al., 2017), on wild-type mice. This vector has been shown to be transported anterogradely down the axon and to enable expression of the Cre-recombinase in postsynaptic target neurons that then are amenable to visualization through Cre-dependent transgene expression. We therefore injected, one week after the first viral injection in the spinal cord, a Cre-dependent AAV-eYFP vector in the pF region. This led to numerous eYFP-expressing cells (Figure 4h). Since this region does not project to the lumbar spinal cord (Figure S6), the Cre-dependent labelling cannot owe to spurious retrograde transport of the AAV1-Syn-Cre virus. Therefore, ascending spinal projections synaptically target the pF respiratory region.

**Figure 4:**
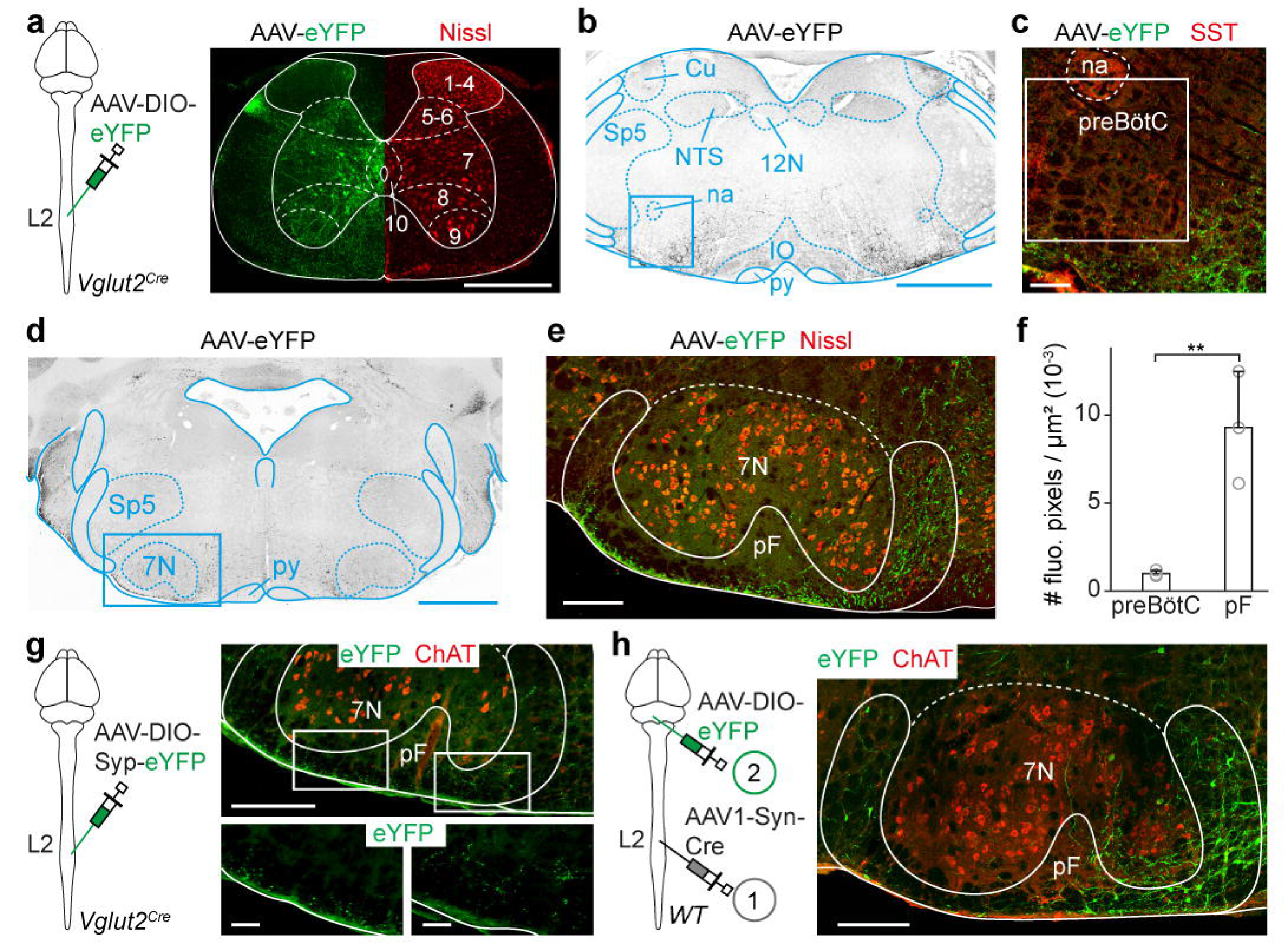
Glutamatergic neurons of the lumbar spinal cord contact the pF respiratory area. **(a)** Experimental strategy for labelling the projections of glutamatergic lumbar neurons: an AAV-DIO-eYFP is injected bilaterally in the 2^nd^ lumbar spinal segment (L2) in a *Vglut2^Cre^* adult mouse. Transfected cells are concentrated in ventral laminae (delineated using a Nissl counterstain) except lamina 9 containing motoneurons. Scale bar, 500 µm. **(b)** Transverse brainstem section at the level of the preBötC showing eYFP^+^ projections in grey. Scale bar, 1 mm. **(c)** Magnification of the blue boxed area that depicts the 400 x 400 µm square used to delineate anatomically the preBötC in this study, with a counterstaining for somatostatin (SST). Note the absence of eYFP^+^ projections. Scale bar, 100 µm. **(d)** Transverse brainstem section at the level of the pF showing eYFP^+^ projections in grey. Scale bar, 1 mm. **(e)** Magnification of the boxed area. The pF area is framed in white. Note the abundant eYFP^+^ projections from the spinal cord. (**f**) Bar-graphs showing the mean density ± SD across mice of eYFP^+^ fluorescent pixels in the preBötC and the pF (both sides pooled). Open gray circles are the mean values of individual mice. **p < 0.01 (Wilcoxon matched-pairs tests; 9 sections from n = 3 mice). **(g)** Bilateral injection, in the L2 segment of a *Vglut2^Cre^* adult mouse of an AAV-DIO-Syp-eYFP. Labelled boutons are detected in the pF on transverse sections, ascertaining that ascending projections from the spinal cord form synaptic contacts. Scale bars, 250 (top) and 50 (bottom) µm. Representative of n = 3 mice. **(h)** Anterograde transynaptic tracing from the ventral lumbar spinal cord using an AAV1-Syn-Cre injected bilaterally at L2 followed by an AAV-DIO-eYFP injected in the pF area in a wild-type adult mouse. The transverse section shows eYFP^+^ neurons in the pF area, further confirming that spinal ascending projections establish synaptic contacts in this area. Representative of n = 3 mice. Scale bar, 200 µm. See also Figure S6.

### Lumbar locomotor circuits upregulate breathing rate through the RTN*^Phox2b/Atoh1^ ex vivo*

To investigate functionally the possibility that lumbar locomotor circuits can upregulate breathing, we used the *ex vivo* isolated brainstem/spinal cord preparation from neonatal mouse. Although long-used for monitoring locomotor (Kjaerulff and Kiehn, 1996; Bouvier et al., 2015) or respiratory-like (Bouvier et al., 2010; Ramanantsoa et al., 2011a) activities, monitoring both simultaneously using pharmacological activation of the lumbar CPG had only been achieved on neonatal rat preparations (Le Gal et al., 2014; Le Gal et al., 2020). We adapted the method to the neonatal mouse using a split-bath allowing independent pharmacological manipulation of the brainstem and spinal cord superfused by dedicated artificial- cerebrospinal fluids (a-CSFs, see methods). In these conditions, we recorded respiratory-like activity on the 4^th^ cervical ventral root and locomotor-like activity on the 2^nd^ lumbar ventral root (Figure 5a, b).

**Figure 5:**
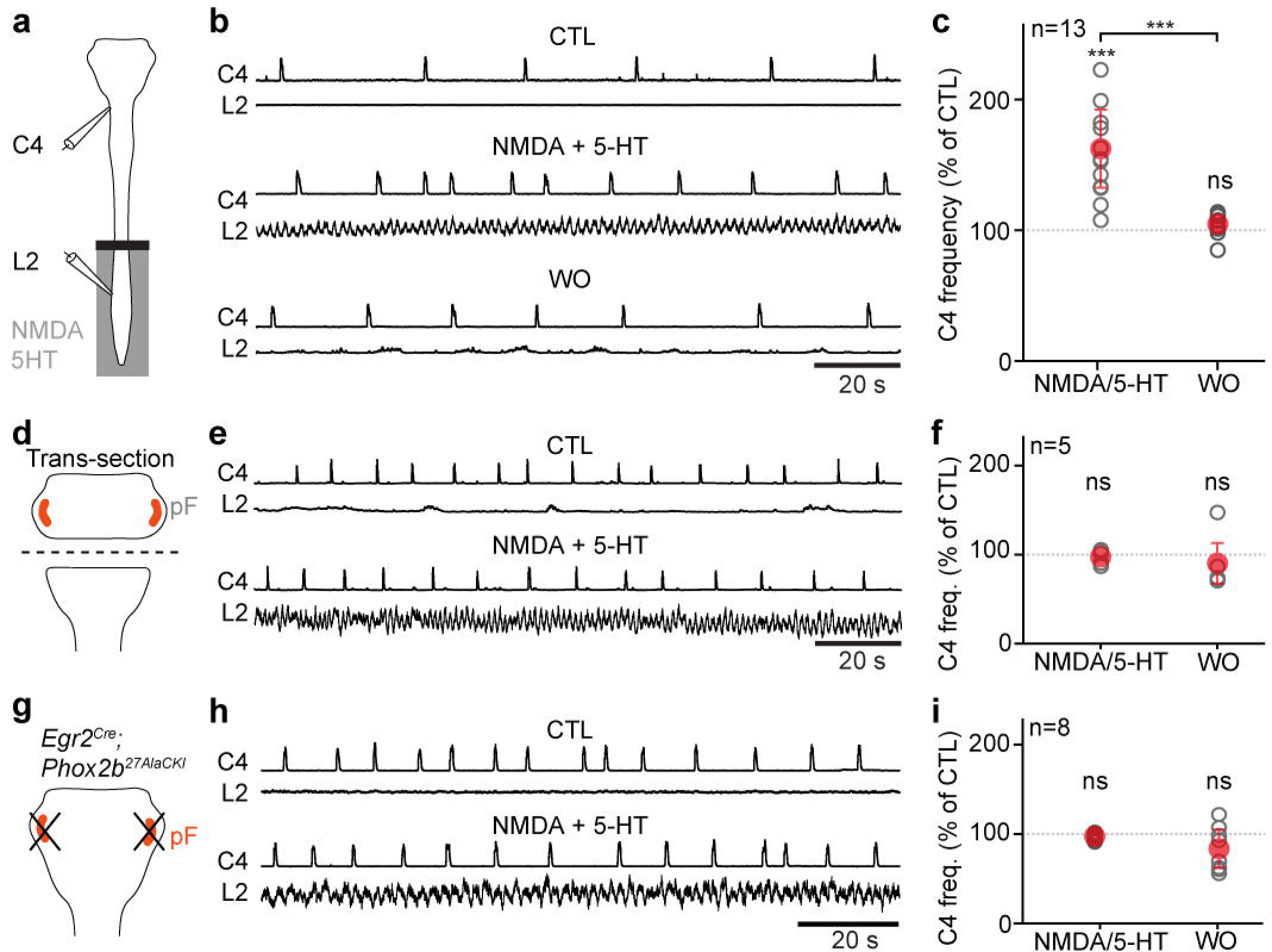
Pharmacological activation of locomotor-like activities increases inspiratory-like frequencies *ex vivo*, which requires the *Atoh1^+^*/*Phox2b^+^* RTN integrity. **(a)** Schematics of the isolated *ex vivo* brainstem-spinal cord preparation from neonates (1-2 days) used throughout the figure. Respiratory- and locomotor-like activities are recorded using glass suction electrodes attached to the 4^th^ cervical (C4) and the 2^nd^ lumbar (L2) motor nerve roots respectively. The preparation is separated in two compartments using a Vaseline barrier (black bar). The lumbar spinal cord is superfused with control or locomotor drugs enriched (NDMA and 5-HT, 10-14 µM each) artificial cerebrospinal fluid (aCSF) while the brainstem compartment remains in control aCSF. **(b)** Recordings of C4 and L2 activities of one representative preparation before (CTL), during, and after (washout, WO) perfusion of locomotor drugs on the lumbar spinal cord (only integrated traces are shown). Note the triggering of locomotor-like activity on L2 and a concomitant increase in the respiratory-like frequency on C4. **(c)** Quantification of the C4 frequency change during drug-induced locomotor-like activity (NMDA/5-HT) and during washout, as a percent change to the CTL condition (100 %, grey dotted line). Grey open circles are the means of individual n preparations, and red circles are the means ± SD across preparations. **(d-f)** Similar experiments as in (**a-c**) but a brainstem transection was performed to physically remove the pF respiratory area. Note the absence of frequency change on the C4 root during drug application. **(g-i)** Similar experiments as in (**a-c**) performed in *Egr2^Cre^;Phox2b^27AlaCKI^* neonates, which lack pF neurons that express Phox2b and are derived from rhombomere 5. This intersection recapitulates the genetically-defined RTN*^Phox2b/Atoh1^*. Note the absence of frequency change on the C4 root during drug application. ***, p < 0.001; ns, not significant (Wilcoxon matched-pairs tests). See also Figures S7 and S8.

When both the brainstem and spinal cord were superfused with their respective control a-CSF solution, the frequency of respiratory-like activities was found to be ranging from 2.5 to 8.3 bursts/min, with an average of 4.3 ± 1.6. Bath-application of the neuroactive substances N-methyl-D-aspartate (NMDA) and serotonin (5-HT) in the spinal cord compartment evoked locomotor-like activities associated with an increased frequency of the respiratory-like activity (average of 7.2 ± 3.6 bursts/min which represents 162 ± 30 % of baseline, Figure 5b, c). Reminiscent of what we observed in freely running mice (Hérent et al., 2020), we found that respiratory- and locomotor-like activity were not temporally synchronized (data not shown). To rule out drug leakage from the spinal compartment we first verified that this frequency increase of respiratory-like activities was abolished following a cervical transection of the medulla (Figure S7a-c). We also examined respiratory changes following the engagement of locomotor-like activities by targeted optogenetic activations of lumbar excitatory neurons (*Vglut2^Cre^; ChR2^floxed^*), in the absence of NMDA and 5- HT (Hagglund et al., 2010; Hagglund et al., 2013; Le Gal et al., 2020). In this optogenetic paradigm, the frequency of inspiratory activity was increased similarly to the pharmacologically-induced locomotion (170 ± 38 % from baseline, Figure S7d-f). Altogether, these experiments indicate that the lumbar spinal segments containing the locomotor CPG exert an excitatory effect on respiratory activity.

Since our anatomical observations place the pF respiratory region as a candidate target of ascending pathways from the spinal cord (Figure 4), we next addressed the functional contribution of this region. In a first set of experiments, we eliminated physically the pF by a complete transection below the facial motor nucleus (Figure 5d). Respiratory-like activities persisted on the 4^th^ cervical root, but pharmacological activation of lumbar circuits was no longer capable of significantly upregulating their frequency (Figure 5e, f). In a second set of experiments, we examined specifically the contribution of RTN neurons in the pF region (Abbott et al., 2009; Abbott et al., 2011). For this we made use of the fact that RTN neurons relevant for modulating respiratory rhythm generation, at least in the context of central chemoception, are i) best identified by their combined history of expression of the transcription factors *Phox2b* and *Atoh1* during embryogenesis (thereafter RTN*^Phox2b/Atoh1^* neurons; Ramanantsoa et al., 2011a; Ruffault et al., 2015) and ii) can be deleted when expressing a mutated allele of PHOX2B (*Phox2b^27Ala^*) in rhombomeres 3 and 5 (*Egr2^cre^;Phox2b^27AlaCKI^* mutants; Figure S8; Ramanantsoa et al., 2011a; Ruffault et al., 2015). We thus generated RTN mutant pups and recorded respiratory- and locomotor-like activities as above. Preparations showed persistent inspiratory-like activity on the C4 root, but pharmacological activation of lumbar circuits was no longer capable of significantly upregulating its frequency (Figure 5g-i). These experiments altogether highlight the capacity of spinal lumbar circuits to upregulate respiratory-like activities through RTN*^Phox2b/Atoh1^* neurons in the pF respiratory region.

### Silencing RTN*^Phox2b/Atoh1^* neurons reduces respiratory increase during running exercise *in vivo*

The importance of RTN*^Phox2b/Atoh1^* neurons revealed above on *ex vivo* neonatal preparations prompted to address their contribution to respiratory activity during running in behaving adult mice. For this, we used the *Atoh1^FRTCre^;Phox2b^Flpo^* intersectional background, i.e., in which Cre-expression from the *Atoh1* locus is conditional to Flpo expression driven by the *Phox2b* locus (Ruffault et al., 2015), and injected in the pF region bilaterally a Cre-dependent AAV coding the inhibitory DREADD receptor hM4Di (Figure 6a, b). Three weeks later, animals underwent the chronic implantation of diaphragmatic EMG electrodes (Hérent et al., 2020). Inspiratory frequency was then measured before, and 2-3 h after, the administration of the DREADD ligand Clozapine-N-Oxide (CNO) while animals were at rest or running at a set frequency on a motorized treadmill.

**Figure 6:**
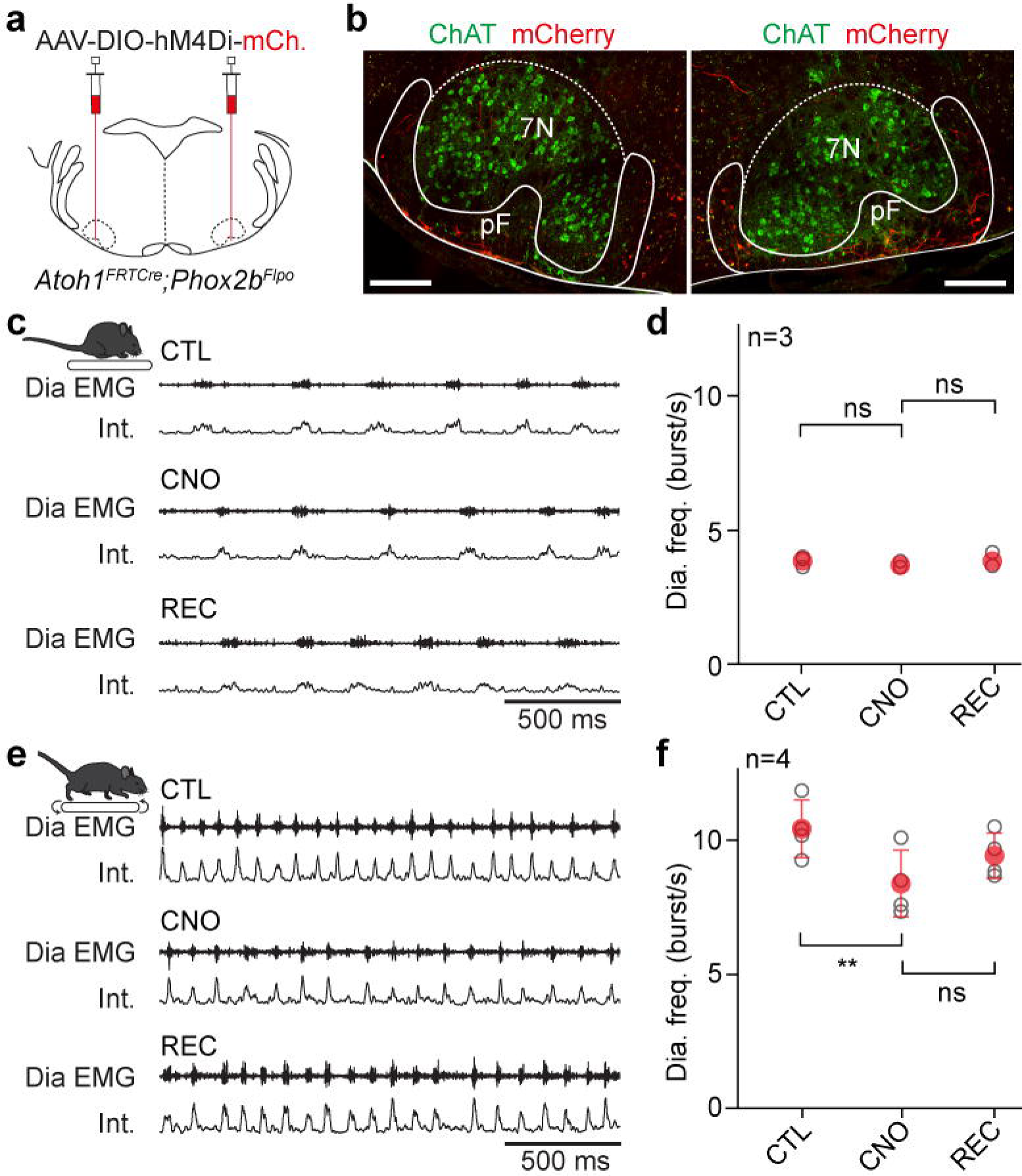
Silencing RTN*^Phox2b/Atoh1^* neurons reduces respiratory frequency during running. **(a)** Experimental strategy for the bilateral transfection of RTN*^Phox2b/Atoh1^* neurons with the inhibitory DREADD receptor hM4Di in an *Atoh1^FRTCre^;Phox2b^Flpo^* adult mouse. **(b)** Example transverse sections showing bilaterally transfected neurons in the pF area ventrally, ventro-laterally, and ventro-medially to the facial motor nucleus (7N) identified by the expression of Choline Acetyl Transferase (ChAT). Scale bar, 250 µm. **(c)** Raw (Dia EMG) and integrated (Int.) recordings of the diaphragm activity of one representative animal at rest before (CTL), during (CNO) and after (REC) intraperitoneal injection of CNO at 10 mg/kg. **(d)** Quantification of the diaphragm mean frequency before, during and after CNO administration at rest. Note that silencing RTN*^Phox2b/Atoh1^* neurons does not alter basal breathing frequency. **(e, f)** Similar representations as in (**c, d**) in mice running on a treadmill at 40 cm/s. Note the significantly lower diaphragm frequency following CNO administration compared to non-injected control mice (CTL). In all graphs, grey open circles are the means of individual mice, and filled circles are the means ± SD across n mice. ns, not significant. **, p < 0.01 (Wilcoxon matched-pairs test in d and and paired t-tests in f). See also Figure S9.

At rest, we found that CNO administration had no significant effect on the mean inspiratory frequency (Figure 6c, d), supporting previous findings that the RTN only minimally contributes to setting the baseline breathing rate (Korsak et al., 2018). When animals were made to run for 1.5 min at a velocity of 40 cm/s set by the treadmill, they presented an augmented respiratory frequency (from 3.9 ± 0.2 bursts/s at rest to 10.4 ± 1.1, which represents a 271 % increase, Figure 6e, f). We have recently reported that this value is characteristic of running mice on a treadmill, regardless of the trotting speed (Hérent et al., 2020). Following CNO administration, while animals were still capable to run at 40 cm/s for 1.5 min, their breathing rate was significantly reduced when compared to controls (8.4 ± 1.2 bursts/s, which represents a 227 % augmentation from rest). Administration of saline in hM4Di-injected mice or of CNO on wild-type mice did not significantly reduce breathing rate during running sessions (Figure S9). These experiments altogether indicate that the activity of RTN*^Phox2b/Atoh1^* neurons is required for setting the adapted ventilatory frequency during running exercise.

### RTN*^Phox2b/Atoh1^* neurons project to the preBötC inspiratory generator

The reduced breathing rate following the silencing of RTN*^Phox2b/Atoh1^* neurons suggests that this genetically defined neuronal subset may have access to the main inspiratory generator, the preBötC. While the pF region was previously shown to send axonal projections to the ventral respiratory column and possibly the preBötC (Smith et al., 1989; Li et al., 2016), this had not been determined for RTN*^Phox2b/Atoh1^* neurons. For this, similarly to what we did for CnF neurons (Figures 1, 2), we injected, in the pF region of *Atoh1^FRTCre^;Phox2b^Flpo^* animals, a Cre-dependent AAV vector coding the ChR2 and a fluorescent protein (AAV-DIO-ChR2-eYFP; Figure 7a, b) and implanted an optic fiber above the injection site. Anatomically, we observed abundant eYFP-labelled varicosities of RTN*^Phox2b/Atoh1^* neurons in the preBötC region with a strong ipsilateral dominance (Figure 7c, d). To demonstrate functionally the capacity of RTN*^Phox2b/Atoh1^* neurons to directly modulate rhythm generation mechanisms in the preBötC, we examined respiratory responses to short single-pulse (50 ms) photoactivations with the same analytic tools described earlier for CnF activations (Figure 2). We found that unilateral photostimulation of RTN*^Phox2b/Atoh1^* neurons could elicit an ectopic inspiratory burst and shorten the respiratory cycle (Figure 7e). However, when compared to CnF photostimulation, shortenings of the respiratory peaked earlier during the inspiration phase (phase 0.2-0.3 and 0.3 - 0.4; phase shift: 0.70 ± 0.13 and 0.69 ± 0.09; p < 0.0001) and, contrary to the CnF, stimulation of the RTN during late expiration caused a significant lengthening of the respiratory cycle (phases 0.9 - 1; phase shift: 1.11 ± 0.12; p = 0.0004; Figure 7f). Therefore here again, stimulation of RTN*^Phox2b/Atoh1^* neurons led to phase dependent inspiratory responses in keeping with the preBötC excitability dynamics. Yet unlike the CnF, no shortening of the subsequent cycles following the stimulus was observed (Figure S10). Moreover, 1 s pulsed-light stimulations of the RTN*^Phox2b/Atoh1^* neurons resulted in significant augmentations of respiratory frequency during the stimulus (Figure 7g-h). The capacity of RTN*^Phox2b/Atoh1^* to contact the preBötC directly is further corroborated by the detection of eYFP puncta in the preBötC following the injection of a Cre-dependent AAV coding a synaptophysin-fused eYFP in the pF area on *Atoh1^FRTCre^;Phox2b^Flpo^* animals (Figure 7i-k). Altogether, these results demonstrate that the genetically defined subset of RTN neurons co-expressing *Atoh1* and *Phox2b* can upregulate respiratory rate through direct projections to the preBötC. Interestingly, the phase-dependent shortening (in inspiration) and lengthening (in late expiration) observed when stimulating RTN*^Phox2b/Atoh1^* neurons are reminiscent of those reported when photostimulating inhibitory preBötC neurons (Baertsch et al., 2018). This suggests that RTN*^Phox2b/Atoh1^* and CnF locomotor-related drives may also differ by their targeting of a distinct balance of excitatory versus inhibitory preBötC neurons.

**Figure 7:**
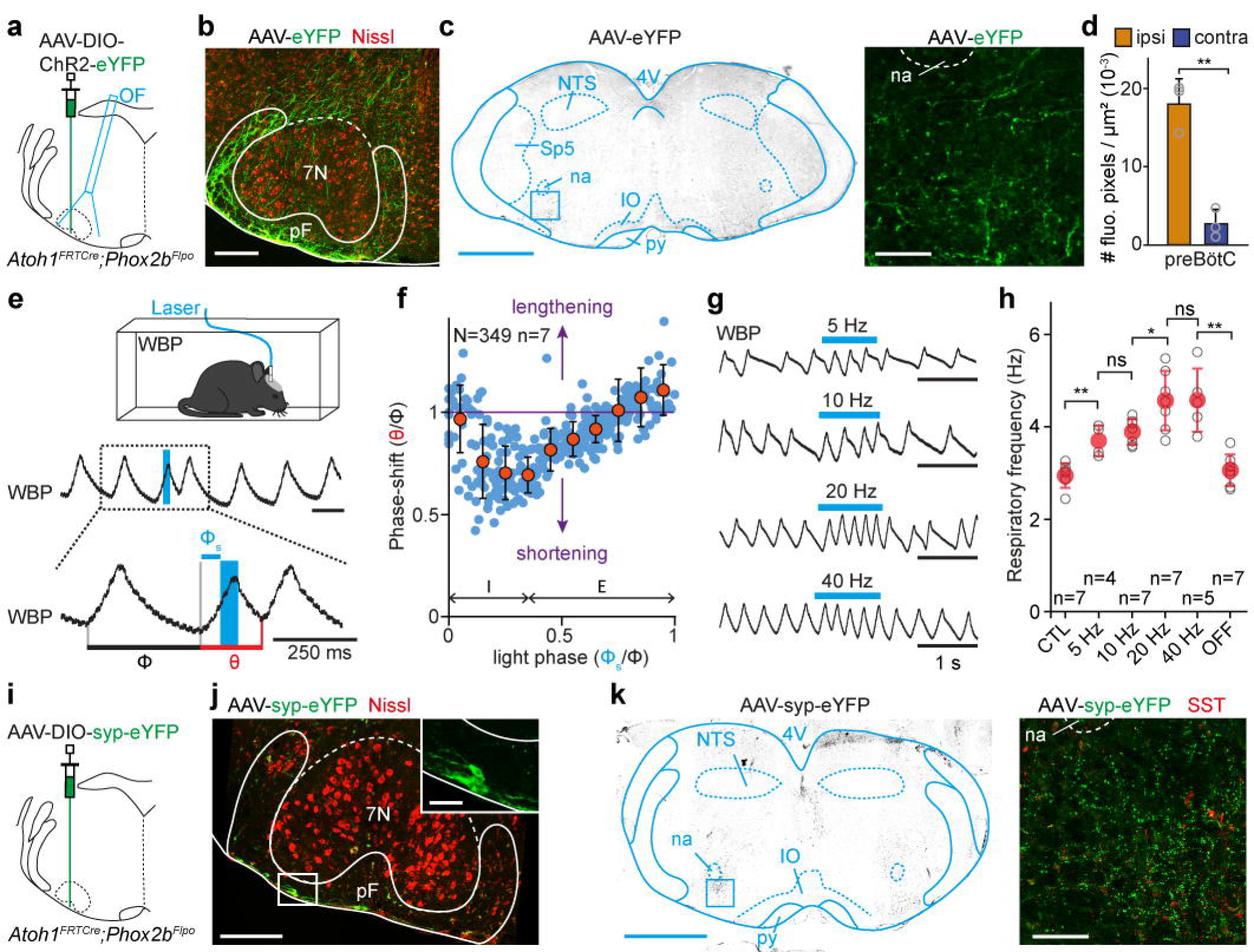
RTN*^Phox2b/Atoh1^* neurons contact the preBötC and impact respiratory rhythm generation. **(a)** Experimental strategy for labelling and photo-activating RTN*^Phox2b/Atoh1^* neurons and projections: an AAV-DIO-ChR2- eYFP is injected in the pF area of an *Atoh1^FRTCre^;Phox2b^Flpo^* adult mouse and an optic fiber (OF) is implanted above the injection site. **(b)** Example transverse section showing unilaterally transfected RTN*^Phox2b/Atoh1^* somata. Scale bar, 250 µm. **(c)** Left: transverse section of the medulla at the segmental level of the preBötC. Scale bar, 500 µm. Right: magnification of the 400 x 400 µm boxed area containing the preBötC, showing abundant eYFP^+^ projections. Scale bar, 100 µm. **(d)** Bar-graphs showing the means density ± SD across mice of eYFP-labelled pixels ipsilaterally and contralaterally in the preBötC. Grey open circles are the mean values of individual mice. **p < 0.01 (Wilcoxon matched-pairs test; 9 sections from n = 3 mice). **(e)** Top: setup for recording ventilation using whole-body plethysmography (WBP) during optogenetic activations. Middle: WBP recordings around a single 50 ms light-stimulation pulse of RTN*^Phox2b/Atoh1^* neurons inspiration (inspirations are upwards, expirations are downwards). Note that the stimulation shortens the respiratory cycle. Bottom: magnification of the above trace showing the control cycle (φ, black), the phase of light-simulation (φs, blue), and the perturbed cycle (θ, red). **(f)** Plot of the phase-shift (perturbed cycle normalized to the control cycle: θ/φ) as a function of the phase of light- stimulation normalized to the control cycle (φs/φ). Note that the phase-shift is respectively shortened (values < 1, purple line) or lengthened (values > 1) when the light pulse occurs in the mid-inspiratory/early-expiratory or late-expiratory part of the respiratory cycle respectively. Inspiration (I) and expiration (E) mean durations are indicated. Blue circles represent individual data from N = 349 random trials from n = 7 mice. Orange circles are averages ± SD across all trials within 0.1 ms bins. **(g)** WBP recordings of ventilation during photoactivation of RTN*^Phox2b/Atoh1^* neurons at increasing train frequencies. **(h)** Quantification of the respiratory frequency before (CTL), during and after (OFF) photostimulations of RTN*^Phox2b/Atoh1^* neurons at different frequencies. Gray open circles are the means of 3 trials for individual animals and filled circles are the means ± SD across n mice. ns, not significant; *, p < 0.05; **, p < 0.01 (Mann-Whitney tests). **(i)** Experimental strategy for labelling the synaptic contacts of RTN*^Phox2b/Atoh1^* neurons using a virally-driven synaptophysin-eYFP fusion protein (Syp-eYFP). **(j)** Example transverse section showing unilaterally transfected RTN*^Phox2b/Atoh1^* somata in the pF area. Scale bar, 250 µm; inset, 50 µm. **(k)** Left: transverse section at the level of the preBötC. Scale bar, 1 mm. Right: magnification of the 400 x 400 µm boxed area containing the preBötC (identified by somatostatin expressing cells, SST), showing abundant eYFP^+^ synaptic contacts of RTN*^Phox2b/Atoh1^* neurons. Scale bar, 100 µm. Representative of n=4 animals. See also Figure S10.

## Discussion

A neuronal substrate for hyperpnea during running has long been proposed but its underlying cells and circuits had remained speculative. We uncover here two systems by which the central locomotor network can enable respiratory rate to be augmented in relation to running activity. On the one hand, we demonstrate the capacity of the MLR subnucleus CnF, a conserved locomotor controller (Shik et al., 1969; Dubuc et al., 2008; Le Ray et al., 2011; Bachmann et al., 2013; Roseberry et al., 2016; Caggiano et al., 2018; Josset et al., 2018; Dautan et al., 2020; Chang et al., 2021; van der Zouwen et al., 2021), to upregulate breathing. On the other hand, we demonstrate that the lumbar enlargement of the spinal cord, containing the hindlimb CPG, also acts as a potent upregulator of breathing rate. Using cell-type specific circuit tracing, we further characterize both locomotor drives by identifying their distinct neuronal targets in the respiratory network.

### Multiple locomotor drives set respiratory rhythm frequency

One intriguing feature we observed with CnF stimulations is that respiratory rate is upregulated before the engagement of effective locomotor movements, or for stimulations intensities that are below the threshold for locomotor initiation (Figure 3). This might be consequent to the direct access of the CnF to the preBötC (Figures 1, 2), while its access to the limb controller requires the crossing of multiple synapses, including onto relay reticulospinal neurons (Jordan et al., 2008; Capelli et al., 2017; Caggiano et al., 2018) that likely accounts for the latencies observed from light onset to locomotor movements (Figure S4c). The regulation of breathing by the CnF in the absence of locomotor movements, revealing the CnF as a *bona fide* respiratory modulator structure, may bear physiological relevance as an anticipatory mechanism to the planned motor action. In the lamprey, spontaneous swimming bouts are preceded by a marked increase in respiratory frequency, an effect that may incriminate the MLR (Gravel et al., 2007; Gariepy et al., 2012). Men informed of an upcoming exercise (Decety et al., 1991; Decety et al., 1993; Thornton et al., 2001; Williamson et al., 2001) or imagining performing an exercise (Krogh and Lindhard, 1913; Tobin et al., 1986; Green et al., 2007), also show increased ventilation and cardiovascular responses, although less drastic than during actual movements, reminiscent of the “pre-loco” phase resulting here from CnF stimulations (Figure 3). Interestingly, the CnF subcomponent of the MLR is associated with escape-like fast regime running, rather than exploratory locomotion (Caggiano et al., 2018; Josset et al., 2018) and may be part of a larger command system for defensive behaviors in the broad sense (Mitchell et al., 1988a; Depoortere et al., 1990; Korte et al., 1992; Ryczko and Dubuc, 2013). It may therefore bear output connectivity allowing to engage a composite response with both cardiovascular (Mitchell et al., 1988b; Korte et al., 1992), respiratory (this study and Gariepy et al., 2012), and if needed, locomotor components. A higher threshold and longer latency for the latter would ensure sufficient priming of relevant autonomic respiratory and cardiovascular controls. Data in the lamprey indicate that locomotor and respiratory centers are not contacted by the same individual MLR neurons with branched collaterals but rather by distinct neurons (Gariepy et al., 2012). Pending an equivalent examination in mice, it is possible that the MLR may, similarly to other descending motor pathways (Sathyamurthy et al., 2020; Usseglio et al., 2020) host projection-defined subsets that each control one trait of a multi-faceted adaptive response.

We also report the existence of a respiratory-modulatory drive from the lumbar spinal segments that contains the hindlimb locomotor circuits (Grillner, 2006; Kiehn, 2016; Grillner and El Manira, 2020). Such an ascending drive, previously suggested in rats (Le Gal et al., 2014; Le Gal et al., 2020) is demonstrated here in mice by local pharmacological (Figure 5) or optogenetic (Figure S7) activations of the lumbar enlargement on reduced preparations *ex vivo.* Indeed, *ex vivo*, the absence of the MLR and of peripheral structures, as well as the experimental control of the extracellular solution, allow to isolate the functional contribution of the spinal ascending drive from descending, peripheral feedbacks and central chemoceptive ones. Since the activity of the spinal locomotor circuits is proportional to locomotor speed (Talpalar and Kiehn, 2010), it is possible that this ascending drive continuously informs supra-spinal structures including respiratory centers on the state of ongoing locomotion. We thus propose that active locomotor executive circuits are, at least partly, causal to the further increase of respiratory rate seen when animals engage in locomotion (i.e., the “loco” phase) following CnF stimulations. In support of this, we show *ex vivo* that fictive respiratory rates are no longer upregulated by engagement of fictive locomotion after elimination of the RTN*^Phox2b/Atoh1^* (Figure 5). However, acute silencing of the RTN*^Phox2b/Atoh1^* neurons *in vivo* reduces, but does not completely prevent, the capacity to accelerate breathing during treadmill running (Figure 6). This could owe to suboptimal efficiency of the chemogenetic inhibition that relies on a double Flpo and Cre recombination strategy and/or variable levels of expression of DREADDS receptors in transfected neurons only partially reducing neuronal activity. In addition, we cannot rule out a persisting activity of CnF neurons during the running exercise (Caggiano et al., 2018). This leaves open the possibility that, during ongoing running, the two locomotor drives, from the CnF and the lumbar spinal cord, may synergize to set the respiratory frequency.

While our results localize the locomotor ascending drive to the lumbar spinal cord, the identity of incriminated neurons will remain to be characterized. Spinal neurons of V2a, V0V or V3 genetic identity as well as Shox2 and Hb9-expresing ones stand as candidates, by virtue of their localization in ventral laminae, their glutamatergic nature and their direct contribution to locomotor rhythm and pattern (Zhang et al., 2008; Dougherty and Kiehn, 2010; Dougherty et al., 2013; Talpalar et al., 2013; Caldeira et al., 2017). Our results also do not speak to the possible involvement of other central locomotor descending pathways (Jordan et al., 2008). However, the absent phasing between respiratory and locomotor drives or movements reported here during CnF-evoked locomotion and that we have previously described during running (Hérent et al., 2020) together with the persistence of a normal ventilatory response to exercise following the removal of peripheral signals in multiple species (Eldridge et al., 1981; Eldridge et al., 1985; Fernandes et al., 1990), make it very unlikely that phasing signals arising from proprioceptive limb afferences or sensory modalities reporting visceral oscillations are important components of the respiratory adaptation to exercise.

### Different respiratory nodes integrate distinct locomotor drives

Another intriguing observation is that the two revealed locomotor drives target different nuclei in the rhythm generating respiratory network. The CnF connects to the preBötC, the main site of inspiratory rhythm generation (Smith et al., 1991; Del Negro et al., 2018), while the lumbar CPG contacts the pF respiratory region, and possibly the RTN*^Phox2b/Atoh1^* previously implicated in central CO2 chemoception (Abbott et al., 2009; Abbott et al., 2011; Ramanantsoa et al., 2011b; Ruffault et al., 2015) that in turn projects onto the preBötC.

For the former, the connectivity is first demonstrated anatomically by the detection of anterogradely- labelled fibers, and a transynaptic labelling approach initiated from glutamatergic preBötC neurons (Figure 1). This connectivity is also supported functionally, by the capacity of CnF neurons or their projections in the preBötC to impact respiratory rhythm mechanisms (Figure 2). In contrast, we found that ascending projections from the lumbar spinal cord were virtually absent in the preBötC but were dense in the pF (Figure 4), in an area compatible with that of the RTN. The synaptic nature of these ascending projections was ascertained using a synaptic labelling (Figure 4g) and an anterograde transynaptic strategy (Figure 4h, Zingg et al., 2017). The functionality of these ascending projections and the identity of their neuronal targets as RTN*^Phox2b/Atoh1^* neurons were evaluated by loss of function experiments *ex vivo* (Figure 5) and *in vivo* (Figure 6). Therefore, while other pF neurons as well as neighboring adrenergic C1 neurons (Abbott et al., 2011; Guyenet et al., 2013) may also be targeted by the locomotor spinal circuits, the RTN*^Phox2b/Atoh1^* subset may be the prominent integrator of the ascending locomotor drive, at least for the setting of respiratory frequency.

We also show that the RTN*^Phox2b/Atoh1^* neurons in turn project to the preBötC (Figure 7). This makes the preBötC inspiratory generator a final integrator of both the descending (from the CnF) and the ascending (from the lumbar spinal cord) locomotor drives and raises the question of the identity of the targeted neurons. Our transynaptic tracing scheme from the preBötC demonstrates that the CnF targets glutamatergic preBötC neurons (Figure 1h-j). Comparatively however, the number of transynaptically labeled neurons in the pF area was much lower (Figure S1d, e). Moreover, brief CnF photostimulations were most effective to shorten the respiratory cycle when delivered during early expiration but failed when delivered during inspiration (Figure 2). This is reminiscent to what was observed when directly stimulating glutamatergic preBötC neurons collectively (Baertsch et al., 2018) or the *Dbx1*-expressing V0 subset (Cui et al., 2016), the main rhythmogenic candidates (Bouvier et al., 2010; Gray et al., 2010). In contrast, when stimulating the RTN*^Phox2b/Atoh1^*, the shortening of the respiratory cycle is most efficient during inspiration and a significant lengthening of the respiratory cycle is observed when photoactivations are delivered in late expiration (compare Figure 2c with Figure 7f). This permissive action in inspiration and the lengthening in expiration recalls what others have reported when specifically activating inhibitory, but not excitatory, preBötC neurons (Baertsch et al., 2018). The RTN*^Phox2b/Atoh1^* and the CnF might thus preferentially target different cell-types in the preBötC: inhibitory neurons for the former, and glutamatergic ones for the latter. Note that such a bias towards preBötC inhibitory target neurons for the RTN*^Phox2b/Atoh1^* compared to the CnF could would be mechanistically compatible with switching from CnF-induced “pre-loco” moderate respiratory frequency increase, to the higher “loco” respiratory frequency range associated to actual locomotion. Indeed, the current model ascribes a limited ability of preBötC excitatory neurons compared to inhibitory neurons at entraining high frequency rhythms (Baertsch et al., 2018). Although our data are compatible with such a working model, the proposed connectivity will need to be investigated directly by future work.

## Methods

### Mice

C57BL/6J wild-type mice were obtained by Janvier Labs (Le Genest-Saint-Isle, France). *VGlut2-IRES- Cre* animals (therefafter *Vglut2^Cre^*, Vong et al., 2011) and *Ai32(RCL-ChR2(H134R)/EYFP* (thereafter *ChR2^floxed^,* Madisen et al., 2012) were obtained from Jackson Laboratories. To manipulate RTN*^Phox2b/Atoh1^* neurons we used the following mouse lines: *Egr2^Cre^* (Voiculescu et al., 2000) crossed with *Phox2b^27AlaCK^*^I^ (Ramanantsoa et al., 2011a) and *Atoh1^FRTCre^;Phox2b^Flpo^* (Ruffault et al., 2015) mouse lines. Animals were group-housed with free access to food and water in controlled temperature conditions and exposed to a conventional 12-h light/dark cycle. Experiments were performed on animals of either sex, aged 2 to 3 months at the time of first injection. All procedures were approved by the French Ethical Committee (authorization 2020- 022410231878) and conducted in accordance with EU Directive 2010/63/EU. All efforts were made to reduce animal suffering and minimize the number of animals.

### Viruses used

For anterograde tracing and photostimulation of the CnF and its projection in the preBötC, we used a Cre-dependent AAV9-Ef1a-DIO-hChR2(E123T/T159C)-eYFP (Addgene #35509, titer 7.7e12 vp/ml, Mattis et al., 2011) unilaterally in the CnF (50 to 100 nL). For anterograde tracing from the lumbar spinal cord, the same virus was injected bilaterally (600-750 nL each side) in the lumbar segment, and for RTN*^Phox2b/Atoh1^* photostimulation, the injection was unilateral (350-500 nL). For reversible silencing of RTN*^Phox2b/Atoh1^* neurons we used bilateral injections (350-500 nL each side) of an AAV8.2-hEF1a-DIO-hM4Di-mCherry-WPRE obtained from Dr. Rachael Neve (Gene Delivery Technology Core, Massachusetts General Hospital, USA, titer: 5.6e12 vp/ml). For transynaptic labelling of inputs onto preBötC neurons, we used 500 nL of a HSV1-hEF1a-LS1L- TVA950-T2A-rabiesOG-IRES-mCherry obtained from Dr. Rachael Neve (Gene Delivery Technology Core, Massachusetts General Hospital, USA), and 200 nL of EnvA-ΔG-rabies-GFP obtained by the GT3 core (Salk Institute, USA, titer: 9.59e9 vp/ml). For anterograde transynaptic tracing, we used injection of the AAV2/1- hSyn-Cre-WPRE-hGH (UPenn Vector Core, titer: 6.68e13 vg/ml, Zingg et al., 2017) bilaterally in the spinal cord (600-750 nL each side) and AAV-DJ-EF1-DIO-hChR2(E123T/T159C)-p2A-eYFP-WPRE (Addgene # 35509, titer: 0.78e12 vg/ml) bilaterally in the pF area (300 nL). For anterograde synaptic tracing from the RTN*^Phox2b/Atoh1^* we used a unilateral injection (350-500 nL) of an AAV8.2-hEF1a-DIO-synaptophysin-eYFP obtained from Dr. Rachael Neve (Gene Delivery Technology Core, Massachusetts General Hospital, USA, titer: 5.4e12 vp/ml).

### Surgical procedures

#### Injections and implants in the brainstem

Animals were anesthetized with isoflurane throughout the surgery (4 % at 1 L/min for induction and 2-3 % at 0.2 L/min for maintenance). Buprenorphine (0,025 mg/kg) was administered subcutaneously for analgesia before the surgery. The temperature of the mice was maintained at 36 °C with a feedback- controlled heating pad. Anesthetized animals were placed on a stereotaxic frame (Kopf) and the skull was exposed. Viral vectors were delivered using a pulled glass pipette connected to a syringe pump (Legato 130, KD Scientific, customized by Phymep, France). The infusion flow was set to 100 nL/min. Coordinates (in mm) used to target CnF neurons (Figures 1-3) were: 4.4 caudal to bregma, 1.3 lateral, and 2.8 mm from the skull surface. RTN*^Phox2b/Atoh1^* neurons were targeted unilaterally (Figure 7) or bilaterally (Figures 4, 6) using the following coordinates: 6.0 caudal to bregma, 1.25 lateral, and 200 µm above the ventral surface. Sequential bilateral injection in the preBötC (Figure 1) were performed at the following coordinates: -7.2 from bregma, 1.25 lateral, and 500 µm above the ventral surface. After the injection, the pipette was held in place for 5 to 10 min before being slowly retracted. For optogenetic activations of the CnF, the CnF fibers in the preBötC and the RTN*^Phox2b/Atoh1^*, a 200 µm core 0.39 NA optic fiber connected to a 1.25 mm diameter ferrule (Thorlabs) was implanted 0.4 mm above the injected site (CnF, Figures 2-3) or 0.8 mm (RTN*^Phox2b/Atoh1^*, Figure 7) or 1.5 mm (preBötC, Figure 2) above the ventral surface. Optic fibers were secured to the skull with dental cement (Tetric Evoflow). Animals were followed daily after the surgery.

#### Injections in the spinal cord

Animals were anesthetized as described above and spinal injections were performed as previously done (Bouvier et al., 2015; Usseglio et al., 2020). A two cm incision of the skin was performed dorsally on anesthetized animals and the exposed spinal column was fixed with two holders on the left and right sides to a stereotaxic frame to minimize movements. Vertebral spinous processes were used as landmarks to target specific segments (Harrison et al., 2013). A small incision of the ligamentum Flavum allowed access to the spinal cord. A pulled glass pipette connected to a motorized syringe pump injector (Legato 130, KD Scientific, customized by Phymep, France) was positioned into the ventromedial area of the L2 (between the 11^th^ and 12^th^ vertebral body) using the following coordinates: 350 μm laterally from the dorsal artery and 800 μm depth from the dorsal surface. This lateral positioning ensures that the injection pipette does not pass through the lateral funiculus where ascending and descending axons travel. For Cholera Toxin B (CTB) experiments (Figure S6), we injected on each side of the spinal cord 600-750 nL of CTB-AF647 conjugate (ThermoFisher Scientific Reference C-34778) diluted at 0.5 % in sterile water. After each injection, the pipette was held in place for 5 to 10 min before being slowly retracted. The skin was sutured, and animals were followed daily after the surgery. All animals recovered without motor impairments.

#### Diaphragm EMG recordings

The protocol was described previously (Hérent et al., 2020). In brief, a 12 cm pair of electrodes was prepared from Teflon-coated insulated steel wires with an outside diameter of 0.14 mm (A-M systems, # 793200). Two wires were lightly twisted together, and a knot was placed 5 cm from one end. At 1 cm from the knot, the Teflon insulation was stripped over 1 mm from each wire so that the two bared regions were separated by about 2 mm. The ends of the two wires were soldered to a miniature dissecting pin. The free ends of the electrodes, as well as a 5 cm ground wire, were soldered to a micro connector (Antelec). Nail polish was used to insulate the wires at the connector.

To implant the diaphragm as previously reported (Hérent et al., 2020), animals were anaesthetized, placed in a stereotaxic frame and hydrated by a subcutaneous injection of saline solution (0.9 %). Their temperature was maintained at 36°C with a feedback-controlled heating pad. This step was crucial to ensure post-surgery survival. The skull was exposed and processed to secure the micro connector using dental cement (Tetric Evofow). The ground wire was inserted under the neck’s skin and the twisted electrodes were tunneled towards the right part of the animal guided by a 10 cm silicon tube of 2 mm inner diameter. The animal was then placed in supine position, the peritoneum was opened horizontally under the sternum, extending laterally to the ribs, and the silicon tube containing the electrodes was pulled through the opening. The sternum was clamped and lifted upwards to expose the diaphragm. A piece of stretched sterile parafilm was placed on the upper part of the liver to avoid friction during movement of the animal and to prevent conjunctive tissue formation at the recording sites. The miniature dissecting pin was pushed through the right floating ribs. The pin was then inserted through the sternum, leaving the bare part of the wires in superficial contact with the diaphragm. The position of the electrodes was secured on both sides of the floating ribs and sternum using dental cement. The pin was removed by cutting above the secured wires. The peritoneum and abdominal openings were sutured, and a head bar was placed on the cemented skull to facilitate animal’s handling when connecting and disconnecting EMG cables during behavioral sessions. Buprenorphine (0.025 mg/kg) was administered subcutaneously for analgesia at the end of the surgery and animals were observed daily following the surgery and treated with Buprenorphine if needed.

### Histology

Adult mice were anesthetized with Euthasol Vet (140 mg/kg) and perfused with 4 % paraformaldehyde (PFA) in 1X Phosphate Buffered Saline (PBS). Brains and spinal cords were dissected out and fixed overnight in 4 % PFA at 4 °C. After fixation, tissues were rinsed in 1X PBS. Brain and spinal cord were cryoprotected overnight at 4 °C, respectively in 16 % and 20 % of sucrose in PBS. Tissues were rapidly cryo- embedded in OCT mounting medium and sectioned at 30 µm using a cryostat. Sections were blocked in a solution of 1X Tris Buffered Saline (TBS), 5 % normal donkey serum and 1 % Triton X-100. The primary antibodies, carried out 48 hours at 4 °C, were: goat anti-ChAt (1:500, ref. # AB144P, Merck Millipore), chicken anti-GFP (1:500, ref. # 1020, Aves Labs), rabbit anti-RFP (1:500, ref. # 600-401-379, Rockland), rabbit anti-SST (1:500, ref. # T-4103, BMA Biomedicals), and sheep anti-TH (1:500, ref. # AB1542, Merck Millipore). Primary antibodies were detected after 2 hours of incubation at room temperature with appropriate secondary antibodies coupled to Alexa-Fluor 488, 647, Cy-3 or Cy-5 (1:500, Jackson ImmunoResearch). Sections were counterstained with a fluorescent Nissl stain (NeuroTrace 435/445 blue, ref. # N21479, 1:200 or NeuroTrace 640/660 deep-red, ref. # N21483, 1:1000, Thermo Fisher Scientific) and mounted in Prolong Diamond Antifade Montant (P36970, Thermo Fisher Scientific) or ibidi Mounting Medium (50001, Ibidi). Sections were acquired with a Leica TCS SP8 confocal microscope (NeuroPICT imaging platform of the NeuroPSI Institute) with 10x and 25x objectives.

The preBötC was defined as located ventrally to the cholinergic ChAT^+^ neurons of the nucleus ambiguus (na) where somatostatin^+^ (SST) neurons are detected (Stornetta et al., 2003, Figures 1, 4, 7, corresponding to areas from 7.0 to 7.4 mm caudal to bregma). The pF respiratory region was defined as immediately ventral, ventro-median and ventro-lateral to the facial motor neurons (7N, Guyenet and Mulkey, 2010, Figures 1, 4, 6, 7, corresponding to areas from 6.5 to 5.7 mm caudal to bregma).

### Behavioral experiments

#### Optogenetic activations

Behavioral experiments started 4 to 5 weeks after the viral injection. Implanted animals were connected to a laser source (473 nm DPSS system, LaserGlow Technologies, Toronto, Canada) through a mating sleeve (Thorlabs). The laser was triggered by the output of a National Instruments interface (NI-USB 6211) and the timings of light activations were delivered using the NI MAX software. For CnF or RTN*^Phox2b/Atoh1^* long photostimulation, light was delivered in trains of pulses of 20 ms (5 to 20 Hz) and of 15 ms (30 and 40 Hz) frequency for a duration of 1 s. Each frequency stimulation was repeated three times with several minutes of rest between trials. We used the minimal laser power sufficient to evoke a response, which was measured to be between 5-12 mW at the fiber tip using a power meter (PM100USB with S120C silicon power head, Thorlabs) to restrict photoactivations unilaterally (Stujenske et al., 2015), prevent heat, and exclude an unintentional silencing by over-activation. For randomized short light-pulses, 50 ms light stimulations (50-70 pulses/experiment) were applied randomly in the respiratory cycle.

#### Plethysmography recordings

To analyze the effect of short photoactivations of the CnF and CnF fibers in the preBötC (Figure 2), or the RTN*^Phox2b/Atoh1^* neurons (Figure 7) on burst timing, ChR2-injected animals were placed inside a hermetic whole-body plethysmography (WBP) chamber (Ruffault et al., 2015), customized to allow the passage of the optical patch-cord, four weeks after viral injection. The plethysmography signal was recorded over a period of 10 minutes using a National Instruments Acquisition card (USB-6211) and the LabScribe NI software (iWorxs).

#### Locomotion in a linear runway

Four to five weeks following the injection of the ChR2-expressing virus in the CnF, animals were implanted with a diaphragm EMG as explained previously (Figure 3). One week following EMG implantation, animals were placed in a linear corridor (80 x 10 cm), and familiarized for 1 h/day for 3 days, prior to the experiments. Implanted animals were filmed from the side at 200 fps and 0.5 ms exposure time using a CMOS camera (Jai GO-2400-USB) and images were streamed to a hard disk using the 2^nd^ LOOK software (IO Industries). The start of the EMG recordings was hardware-triggered by the start of the video-recordings using the frame exposure readout of the video camera, so that the two recordings are synchronized. When animals were immobile at one end of the corridor and the respiration was stable, we delivered CnF optogenetic activations with frequencies ranging from 5 to 40 Hz. For each frequency, the stimulation was repeated three times with several minutes of rest between trials.

#### Chemogenetic silencing and treadmill experiment

Three weeks following the injection of the hM4Di virus in the RTN*^Phox2b/Atoh1^*, animals were implanted for diaphragm EMG recordings as explained above (Figure 6). Non-injected C57BL/6J mice were also implanted as controls to test for CNO side effects. One week following EMG implantation, implanted animals were familiarized on a stationary custom-made motorized treadmill with adjustable speed range (Scop Pro, France, belt dimensions: 6 cm x 30 cm) for 30 min/day for 3 days, prior to the experiments. In addition, implanted animals were exercised during this time for a total of 5 min at 40 cm/s each day. Mice could rest for 5 min after running before being placed back in their cage. This step was crucial to obtain stable running animals during experimental sessions. Following this short training, implanted mice were connected with custom light-weight cables to an AC amplifier (BMA-400, CWE Inc.) and neurograms were filtered (high-pass: 100 Hz, low-pass: 10 kHz), collected at 10 kHz using a National Instruments acquisition card (USB-6211) and live-integrated using the LabScribe NI software (iWorxs). Animals were first placed on the stationary treadmill to monitor basal respiration. Animals were then challenged to trot at 40 cm/s for 1.5 min before being administered intraperitoneally with CNO (Enzo Life science, ref. #: BML-NS105-0005, 10 mg/kg) or saline (0.9 %). Animals were placed again on the treadmill with the same paradigm 2-3 h and 5 h after CNO or saline administration to measure respiration in resting and running conditions. During experiments, animals were filmed from the side in the same way as above to monitor the stability of running episodes.

#### Ex vivo brainstem-spinal cord experiments

Pups aged 1-2 days were used in all experiments. The pups were anaesthetized with isoflurane, decerebrated and the brainstem still attached to the spinal cord was dissected and isolated in ice-cold Ringer’s solution that contained (in mM): 111 NaCl, 3 KCl, 25 NaHCO3, 1.25 MgSO4, 1.1 KH2PO4, 2.5 CaCl2 and 11 D-Glucose, and oxygenated in 95 % O2, 5 % CO2 to obtain a pH of 7.4. Isolated brainstem-spinal cords were transferred into a recording chamber and pinned to a Sylgard 184 resin. Preparations were partitioned in two compartments at the level of lower thoracic segments (T11) using a Vaseline wall, to restrict bath application of locomotor drugs on the lumbar spinal cord (Figure 5). The lumbar compartment was continuously perfused with the above Ringer’s solution while the rostral compartment containing the brainstem was superfused with a Ringer’s solution that contained (in mM): 111 NaCl, 8 KCl, 25 NaHCO3, 3.7 MgSO4, 1.1 KH2PO4, 1.25 CaCl2 and 30 D-Glucose. All recordings were done at room temperature (25°C) after allowing 30 min recovery period after the dissection. Respiratory- and locomotor-like activities were recorded respectively on the 4^th^ cervical (C4) and the 2^nd^ lumbar (L2) ventral roots using extracellular suction glass pipettes (120F-10, Harvard Apparatus). Drug-evoked locomotor-like activities were induced by bath-applying 10 to 14 µM of N-methyl-D-aspartate (NMDA, Tocris) and serotonin (5-HT, Sigma-Aldrich, Figure 5) or using blue light on the lumbar spinal cord of ChR2-expressing pups (*Vglut2^Cre^;ChR2^floxed^*, Figure S7). Signals were collected and band-passed filtered at 100 Hz to 1 kHz with an AC amplifier (Model 1700, A-M Systems) and live-integrated (Neurolog NL703, Digitimer) with a time constant of 100 ms (C4) or 200 ms (L2). Signals were sampled using Clampex 11 (Molecular Devices) at 5 kHz. To control for locomotor drugs leakage, some preparations were transected at the level of the cervical spinal cord (Figure S7a-c). For brainstem transection experiments, the rostral part of the brainstem containing the pF respiratory region was physically removed (Figure 5d-f). *Egr2^Cre^;Phox2b^27AlaCKI^* pups were used to genetically eliminate RTN*^Phox2b/Atoh1^* neurons (Ramanantsoa et al., 2011a; Ruffault et al., 2015, Figure 5g-i).

### Quantifications and statistical analysis

#### Phase-shift analysis

We measured the duration of the respiratory cycle containing the light stimulus (perturbed cycle, θ) and the previous respiratory cycle (control cycle, ɸ, Figures 2 and 7). One respiratory cycle was defined as from the onset of inspiration to the subsequent inspiratory onset. The phase of light-stimulation ɸS was defined as from the onset of the perturbed cycle to the onset of the light pulse. The perturbed cycle θ was defined as from the onset of the inspiration that precedes the light stimulation to the onset of the subsequent inspiration. The perturbed phase (phase-shift) was calculated as the ratio of the perturbed cycle divided by the control cycle (θ/ɸ). The light phase was defined as the ratio of the stimulated cycle divided by the control cycle (ɸS/ɸ). The perturbed phase was then plotted against the light phase for all events from all animals. The number of events (N) and animals (n) are given in the corresponding figures for all tested condition. In addition, the average perturbed phase was plotted against the average light phase in 0.1 ms bins as mean ± SD. Inspiratory time (I) was measured, averaged for each animal and a grand average was calculated and annotated in the corresponding figures for all tested condition. Expiratory time (E) was calculated from respiratory cycle and inspiratory (I) times.

#### Locomotor parameters analysis

To track the mouse displacement and measure its speed, we used DeepLabCut (version 2.1.5.2, Mathis et al., 2018) and manually labelled the positions of the head from 50 frames of each video. We then used 95 % of the labelled frames to train the network using a ResNet-50-based neural network with default parameters for 3 training iterations. This network was then used to analyze videos from similar experimental settings. For treadmill experiments, the head X coordinate was used as a control for running stability on the treadmill. For CnF stimulations on the corridor, the head X coordinate was used to calculate the animal’s speed sx using the gradient over time.

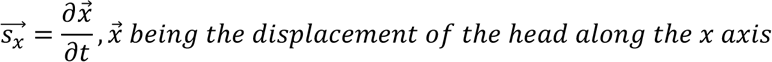

Head X coordinate (treadmill) and calculated velocity (corridor) were then exported to Clampfit (Molecular Devices) and interpolated to 10 kHz to match the acquisition framerate of diaphragmatic EMG recordings. Both sets of signals (head X and diaphragm, or speed and diaphragm) were merged in single files, before being processed offline in Clampfit. The animal’s instantaneous speed is illustrated on figures (Figure 3). The mean speed, defined from the onset of the movement to the end of the photostimulation, was then calculated using the statistic function in Clampfit for each CnF stimulation (5 to 40 Hz). All values were averaged across trials for each animal (3 trials/animal), and a grand mean ± SD across n animals was calculated per stimulation frequency (Figure S4b). Locomotor onset latencies (Figure S4c) were defined as the delay between the onset of the CnF stimulation and the onset of movement for each CnF stimulations. All values were averaged across trials (3 trials/animal) and a grand mean ± SD across n animals was calculated per stimulation frequency.

For gait analysis during CnF photostimulations (Figure S4d), we manually annotated the paw of a reference hindlimb (ipsilateral) and registered the timings of footfalls (when the paw first touches the floor). Each reference locomotor cycle was then defined as the duration from one footfall (ipsi_FFn) to the next (ipsi_FFn+1). The time of occurrence of the contralateral hindlimb footfall within the reference locomotor cycle was annotated manually (contra_FF) and the synchronicity rate was then computed as follows:

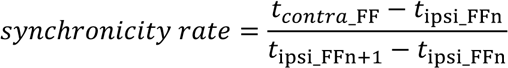

A custom MATLAB script was then used to categorize synchronized (synchronicity rate ∈[0, 0.25[∪[0.75, 1]) or alternated (synchronicity rate ∈[0.25, 0.75[) ipsi- and contralateral hindlimb steps. Synchronicity rates were averaged across animals (3 trials/animal) and a grand mean ± SD across n animals was calculated per stimulation frequency.

#### Locomotor/respiratory coordination

The temporal coordination of breaths to strides (Figure S5) was represented with circular statistics, similarly to what we performed recently (Hérent et al., 2020) and imprinting from numerous studies having investigated the cycle-to-cycle correlations of motor activities (Kjaerulff and Kiehn, 1996; Talpalar et al., 2013; Skarlatou et al., 2020). The phase of each individual inspiratory burst within the locomotor cycle (ΦInsp, around 15 bursts) is represented as the position, from 0 to 1, of the diamond marks on the outer circle (see Figure S5d). For each animal, we also computed the mean phase of consecutive inspiratory bursts and represented it as a colored circle (the mean phases of different animals are in different colors). The distance R of the mean phase to the center of the circle indicates the concentration of individual phase values around the mean, as established by (Kjaerulff and Kiehn, 1996). If inspiratory and locomotor movements are temporally correlated, then individual phase values will be concentrated around a preferred phase value (for instance 0 or 1, at the top of the circle, if the two motor activities were in phase). The mean value would then be positioned at a significant distance from the center. Conversely, if inspiratory and locomotor movements are not coupled, individual phases will be evenly distributed across the circle. Consequently, the mean phase value will be at a short distance from the diagram center, illustrating the dispersion of values around the mean. The inner circles of the circular diagrams depict the threshold for mean phase values to be considered significantly oriented (R < 0.3) as commonly done (Kjaerulff and Kiehn, 1996; Talpalar et al., 2013; Hérent et al., 2020; Skarlatou et al., 2020). Circular plots were obtained using a custom macro in Excel.

#### In vivo respiratory changes analysis

To analyze breathing changes resulting from CnF (EMG recordings, Figure 3) and RTN*^Phox2b/Atoh1^* photostimulation (WBP recordings, Figure 7), instantaneous inspiratory frequencies were detected over a 1 s window, using the threshold search in Clampfit before, during and directly after a 1 s light stimulation for all frequencies (5 to 40 Hz). For CnF stimulations that triggered locomotor episodes (15 to 40 Hz), the recovery period was measured as soon as the animal returned to immobility. Breaths detected from the onset of the light stimulus to the onset of the movement were categorized as during the “pre-loco” (Figure 3d). Breaths detected from the onset of the movement to the offset of the light stimulus were categorized as during the “loco” phase (Figure 3d, f). All values were averaged across animals (3 trials/animal) and a grand mean ± SD across n animals was calculated per stimulation frequency.

For treadmill running (Figure 6), the mean diaphragm frequency was analyzed prior to exercise (resting condition) and from stable trotting moments, i.e., when the animal’s speed was in phase with the treadmill, inferred by the absence of changes in head’s X coordinates (running condition). Instantaneous respiratory frequency was measured for a total duration of 6 s in each condition: before (CTL), during (CNO/saline) and after (REC) administration of either CNO or saline. These measurements were done using 2 to 3 windows taken during resting conditions and at any stable moment of the 1.5 min run (excluding the first 20 seconds to exclude possible stress-induced changes when the treadmill is just engaged). Measurements were averaged to give the mean value for each animal. Averaged mean values were expressed as mean ± SD across n animals.

#### Ex vivo respiratory-like activities analysis

Instantaneous respiratory-like frequencies were analyzed offline using the threshold search in Clampfit (Molecular Devices) before, during and after bath application of NMDA and 5-HT (Figure 5). Respiratory frequency changes during drug and washout conditions were normalized and expressed as a percent of the control. A grand mean ± SD across n animals was calculated.

#### Statistical Analysis

All data are expressed as mean ± SD. Statistical tests were performed using GraphPad Prism 7 and are spelled out in the figure legends, as well as the number of trials (N) and animals (n) used for each experiment. Changes were considered as not significant (ns) when p > 0.05 and as significant when p < 0.05. Significance levels are reported as follows: *, p < 0.05; **, p < 0.01; ***, p < 0.001; and ****, p < 0.0001.

## Acknowledgements

This work was funded by Agence Nationale de la Recherche (ANR-17-CE16-0027 and ANR-20-CE16- 0026 to JB and ANR-15-CE16-0013 to GF) and CNRS, Université Paris-Saclay and NeuroPSI. CH held a doctoral fellowship from Région Ile-de-France and a fellowship extension from Fondation pour la Recherche Médicale (contract # FDT201904007982). We thank Aurélie Heuzé for lab management and genotyping animals, Edwin Gatier for help with DeepLabCut and analytic scripts, Jean-François Brunet for providing the *Atoh1^FRTCre^;Phox2b^Flpo^* animals, and the animal facility for housing animals.

## Author contribution

J.B. and G.F. designed the study and provided funding. J.B. supervised the work. C.H. performed and analyzed experiments with contributions from S.D. and J.B. J.B. and C.H. prepared figures. J.B. wrote the paper and all authors contributed to its editing.

