## Supplemental Figures and Legends for "Upregulation of breathing rate during running exercise by central locomotor circuits"

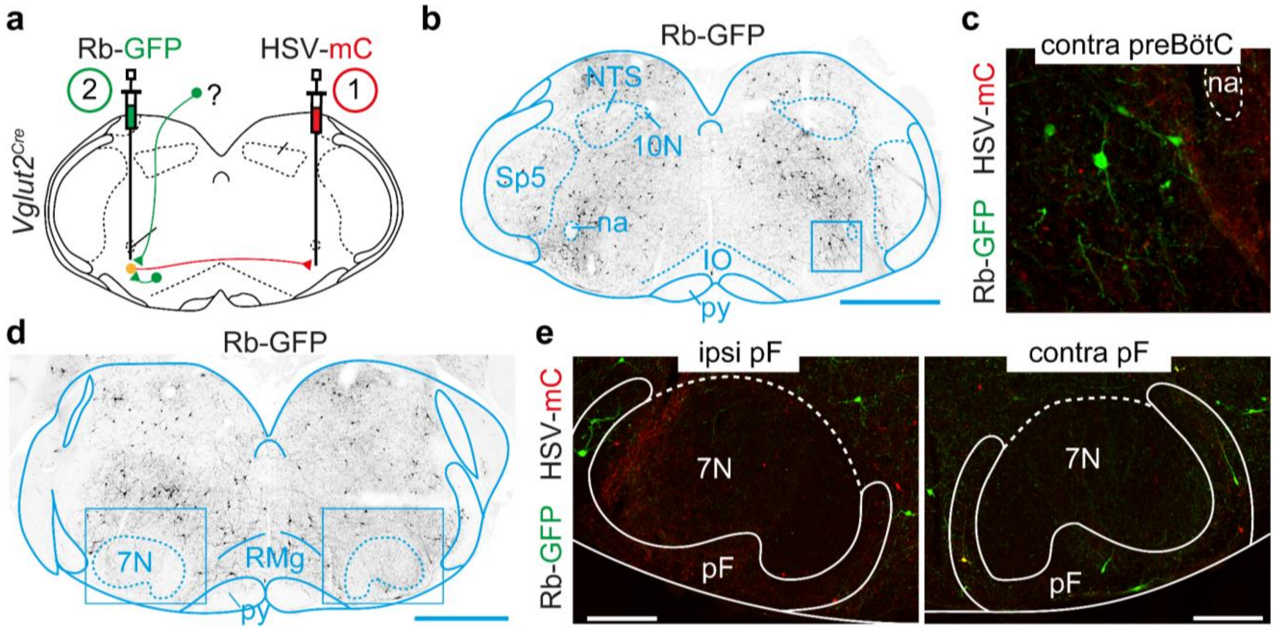


**Figure S1: Transynaptic labelling from Glut+ commissural preBötC neurons leads only little labelling in the pF. Related to Figure 1.**

**(a)** Retrograde transynaptic monosynaptic tracing from glutamatergic commissural preBötC neurons. An HSV-LS1L-TVA-oG-mCherry is first injected in the contralateral preBötC followed by a G-deleted and EnvA pseudotyped Rb virus (EnvA-ΔG-Rb-GFP) in the ipsilateral preBötC in a *Vglut2*$^{Cre}$ adult mouse.

**(b)** Transverse section of the caudal medulla at the preBötC level showing putative presynaptic cells (Rb-GFP$^{+}$). Scale bar, 1 mm.

**(c)** Enlargement of the 400 x 400 µm boxed area containing the preBötC showing the presence of transynaptically labelled cells contralaterally. Scale bar, 100 µm.

**(d)** Transverse section in the rostral medulla at the level of the pF showing the scarcity of presynaptic cells (Rb-GFP$^{+}$). Scale bar, 1 mm.

**(e)** Enlargement of the boxed areas over the ipsilateral and contralateral pF areas. Scale bars, 250 µm.

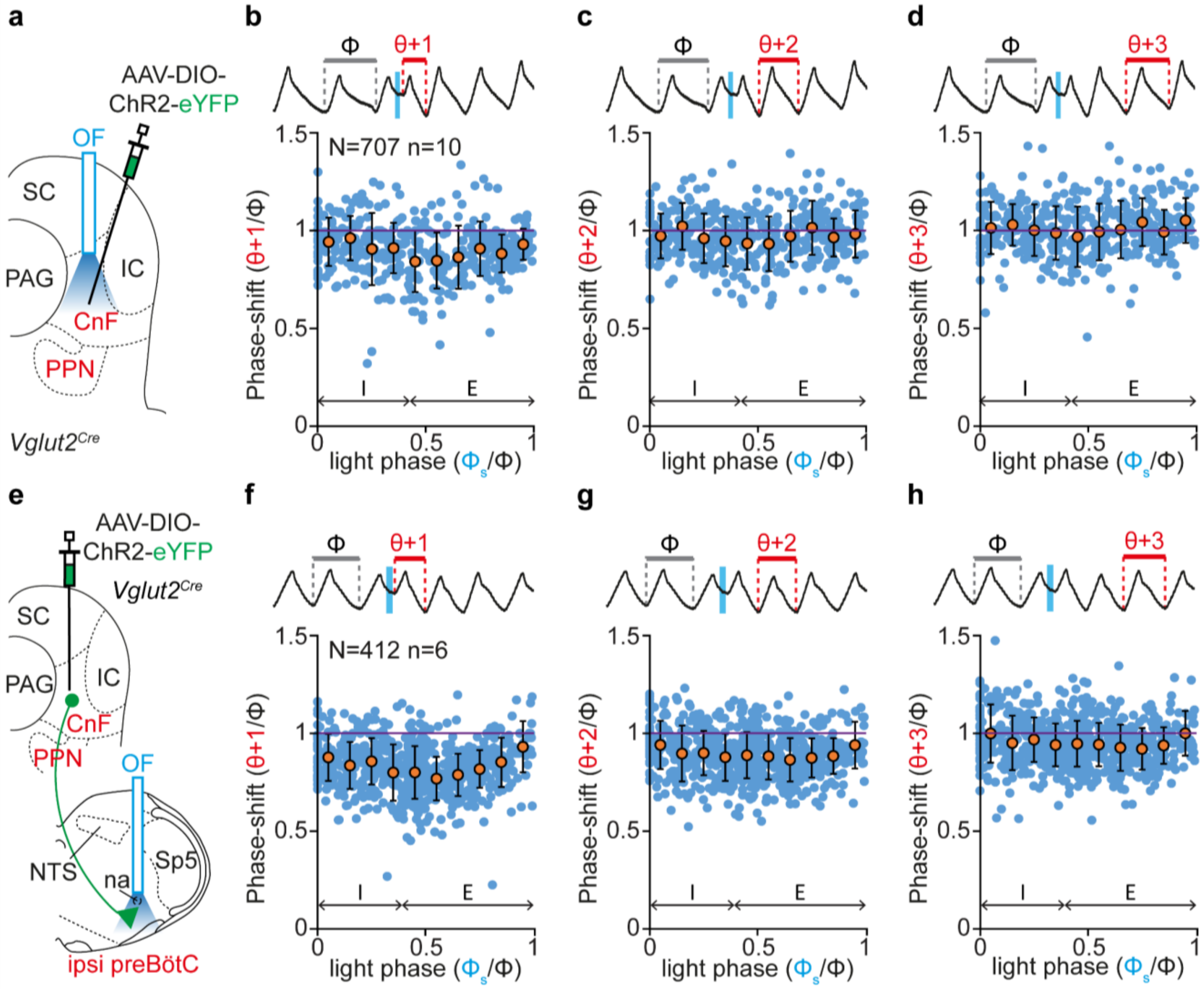


**Figure S2: Impact of photoactivating glutamatergic CnF neurons and their projection to the preBötC area on respiratory cycles following the light-perturbed cycle. Related to Figure 2.**

**(a)** Experimental strategy for photoactivating the glutamatergic CnF neurons: an AAV-DIO-ChR2-eYFP is injected in the CnF of a *Vglut2$^{Cre}$* mouse line and an optic fiber (OF) is implanted above the injection site.

**(b)** Top: whole body plethysmography (WBP) recordings of respiratory cycles while a 50 ms pulse stimulation is applied (inspirations are upwards, expirations are downwards). The cycle n+1 following the perturbed cycle is annotated on the trace (θ+1). Bottom: plot of the perturbed phase+1 (perturbed cycle n+1 normalized to the control cycle: θ+1/ϕ) as a function of the light phase (light cycle normalized to the control cycle: $ϕ_s/ϕ$). Values < 1 (purple line) indicate a shortening of the perturbed cycle. Blue circles represent individual data from N = 707 random trials from n = 10 mice and orange circles are averages ± SD across all trials within 0.1 ms bins. Inspiration (I) and expiration (E) mean durations are annotated on the graph.

**(c, d)** Similar representations for cycles n+2 (θ+2, in c) and n+3 (θ+3, in d).

**(e)** Experimental strategy for photoactivating the fibers of glutamatergic CnF neurons in the preBötC: an AAV-DIO-ChR2-eYFP is injected in the CnF of a *Vglut2$^{Cre}$* mouse line and an optic fiber (OF) is implanted above the preBötC on the same side.

**(f-h)** Similar representations as in (**b-d**) when activating glutamatergic CnF fibers in the preBötC. N = 412 random trials from n = 6 mice.

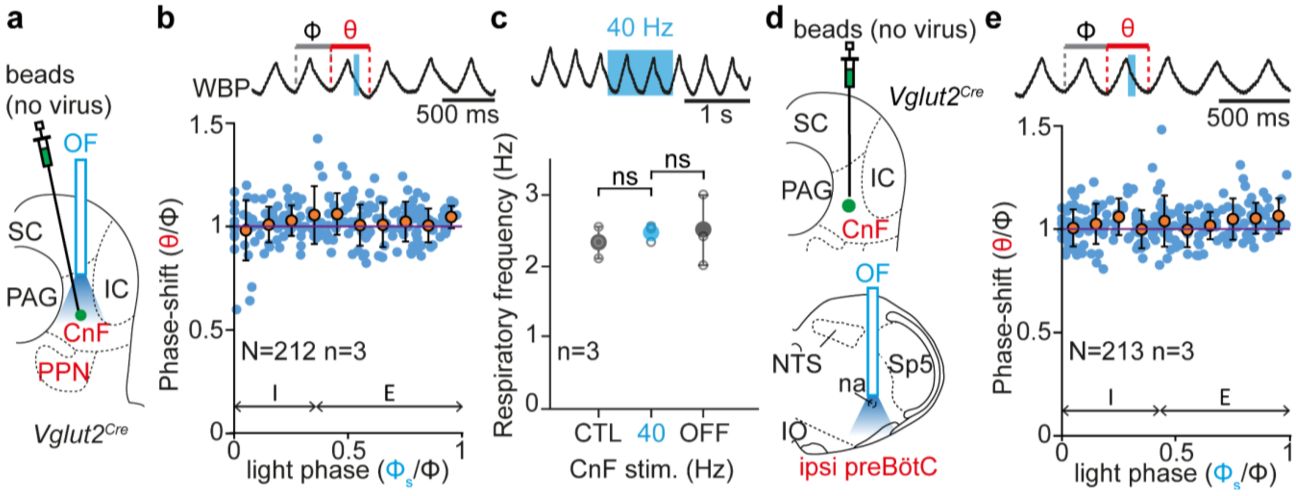


**Figure S3: Controls for optogenetic activations of glutamatergic CnF neurons. Related to Figures 2 and 3.**

**(a)** *Vglut2[cre]* adult mice were injected with fluorescent beads in the CnF, but no ChR2-coding virus. An optic fiber (OF) was implanted above the injection site.

**(b)** Top: Whole-body-plethysmography (WBP) recordings around a single 50 ms light pulse delivered on control mice during the expiratory phase of one respiratory cycle (inspirations are upwards, expirations are downwards). The control cycle (ϕ, black), the phase of light-simulation ($\phi_s$, blue), and the perturbed cycle (θ, red) are indicated (see Figure 2 for details). Bottom: plot of the phase-shift (perturbed cycle normalized to the control cycle: θ/ϕ) as a function of the phase of light-stimulation normalized to the control cycle ($\phi_s/\phi$). Inspiration (I) and expiration (E) mean durations are indicated. Note that, contrary to actual light activations (see Figure 2), the respiratory cycle is not affected by light deliveries on these control mice (values close to 1, purple line). Blue circles represent individual data from N random trials from n mice. Orange circles are averages ± SD across all trials within 0.1 ms bins.

**(c)** Top: example WHB recording during a 1 s light delivery (40 Hz train). Bottom: quantification of the diaphragm frequency showing no significant effect of the light delivery (CTL: control; OFF: after light offset). Gray open circles are the means of individual animals and colored circles are the means ± SD across n mice. ns, not significant (Wilcoxon matched-pairs tests).

**(d)** *Vglut2[cre]* adult mice were injected with fluorescent beads in the CnF, but no ChR2-coding virus. An optic fiber (OF) was implanted above the preBötC on the same side.

**(e)** Same as in (b) during a single 50 ms light delivery in the preBötC area showing again that, contrary to actual light activations (see Figure 2), the respiratory cycle is not affected by light deliveries on these control mice (values close to 1, purple line).

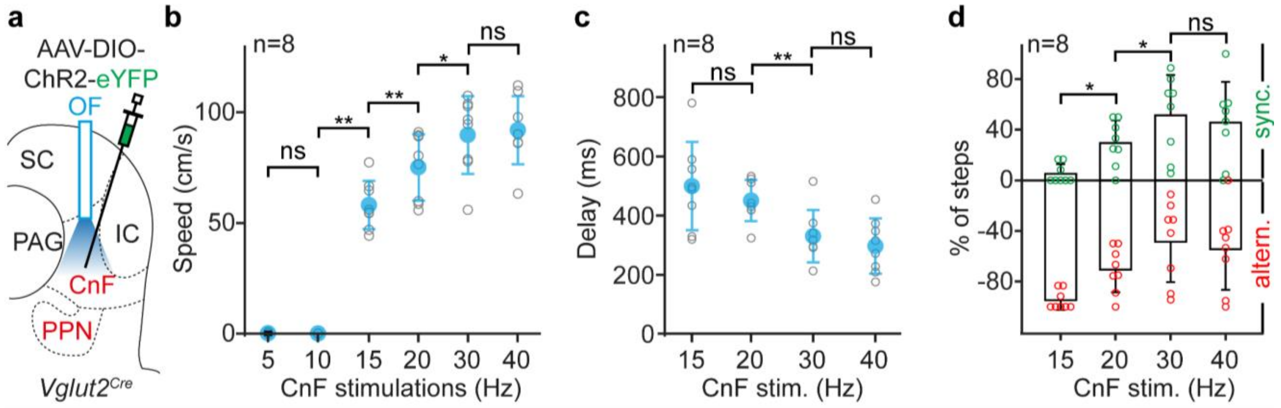


**Figure S4: Glutamatergic CnF neurons control locomotor speed, the locomotor onset delay and the hindlimb synchronicity rate. Related to Figure 3.**

**(a)** Schematics of the viral injection (AAV-DIO-ChR2-eYFP) and of the optic fiber implantation (OF) to photoactivate glutamatergic CnF neurons in an adult *Vglut2$^{Cre}$* mouse.

**(b, c)** Quantification of the animal's displacement speed **(b)** and delay to locomotor initiation **(c)** for increasing CnF photostimulations frequencies. Gray open circles are the means of individual animals and colored circles are the means ± SD across n mice.

**(d)** Quantification of the percent of hindlimb movements that are showing left-right alternation (negative values, indicative of trot) or left-right synchronicity (positive values, indicating of gallop) at increasing CnF photostimulation frequencies. Colored open circles are the means of individual animals and bar-graphs are the means ± SD across n mice. *$p < 0.05$; **$p < 0.01$; ns, not significant (Wilcoxon matched-pairs tests).

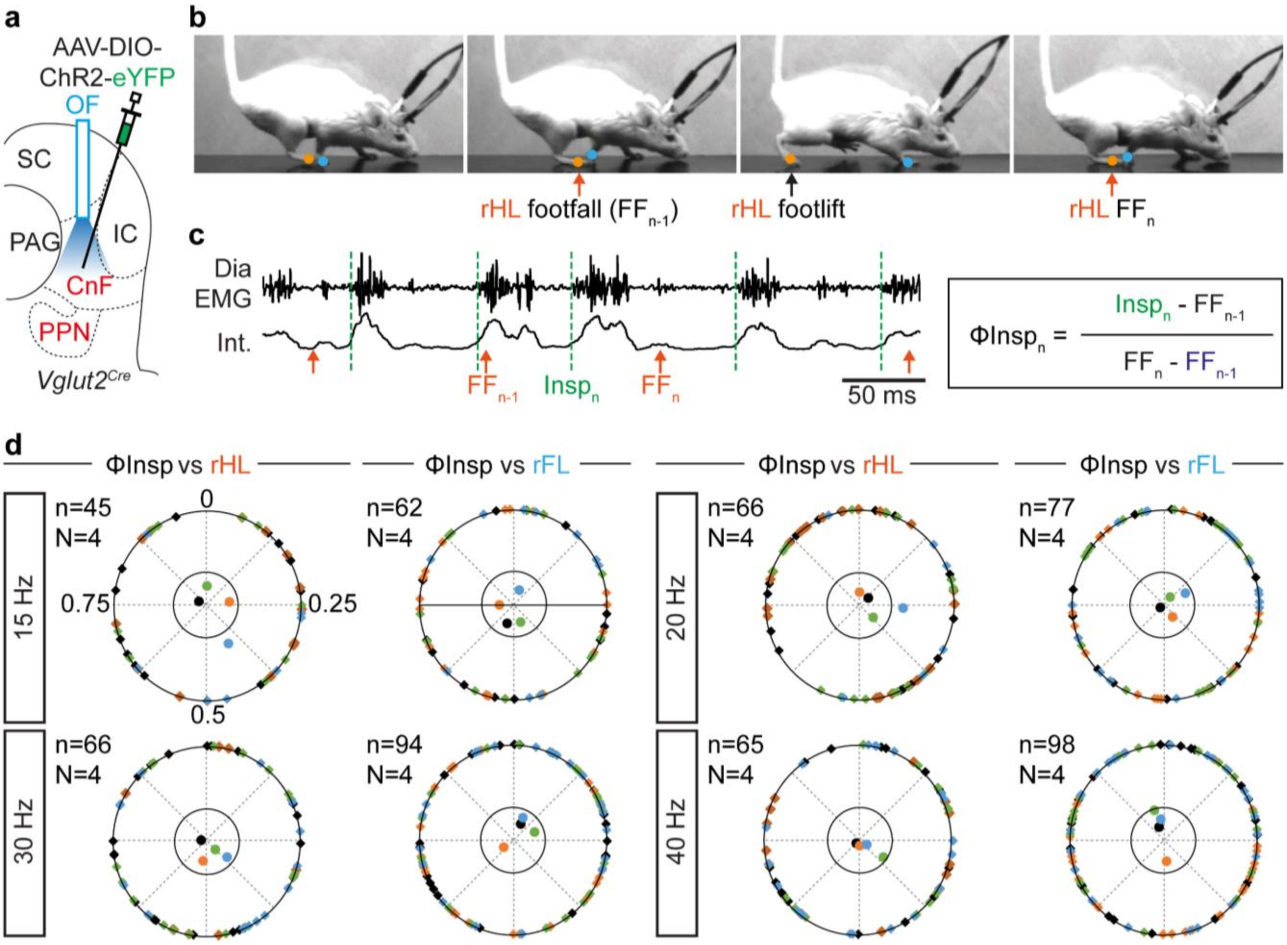


**Figure S5: Absence of phasing of respiratory and locomotor rhythms during CnF-evoked running. Related to Figures 2 and 3.**

**(a)** Schematics of the viral injection (AAV-DIO-ChR2-eYFP) and optic fiber implantation (OF) to photoactivate glutamatergic CnF neurons in an adult *Vglut2$^{Cre}$* mouse.

**(b)** Side views of one representative mouse during CnF photoactivation (20 Hz) on a linear corridor. The timing of footlifts and footfalls of the right hindlimb (rHL, orange) and the right forelimb (rFL, blue) were detected manually on the videos. One complete rHL locomotor cycle is shown, between two consecutive footfalls ($FF_{n-1}$ and $FF_n$).

**(c)** Raw (EMG) and integrated (Int.) diaphragmatic electromyograms and occurrence of rHL footfalls (orange arrows) during CnF-evoked running. Dotted green lines indicate the onsets of inspirations, which can occur at any moment of the locomotor cycle. The occurrences of inspiratory bursts ($Insp_n$) within the locomotor cycle are then calculated and expressed as a phase value ($\Phi Insp_n$) from 0 (concomitant with $FF_{n-1}$) to 1 (concomittant with $FF_n$).

**(d)** Circular plots diagrams showing the phase-relationship between individual inspiratory bursts and the indicated reference limb for one representative animal in response to photoactivations of the CnF at 15, 20, 30 and 40 Hz. Diamonds on the outer circle indicate the phase of n individual inspirations pooled from N mice; values from each animal are given in the same color. The dots inside the circle indicate the mean orientation vector for each animal. The positioning of these mean values within the inner circle illustrates the absence of a significantly oriented phase preference ($R < 0.3$, with R being the concentration of phase values around the mean as defined in Kjaerulff and Kiehn (1996).

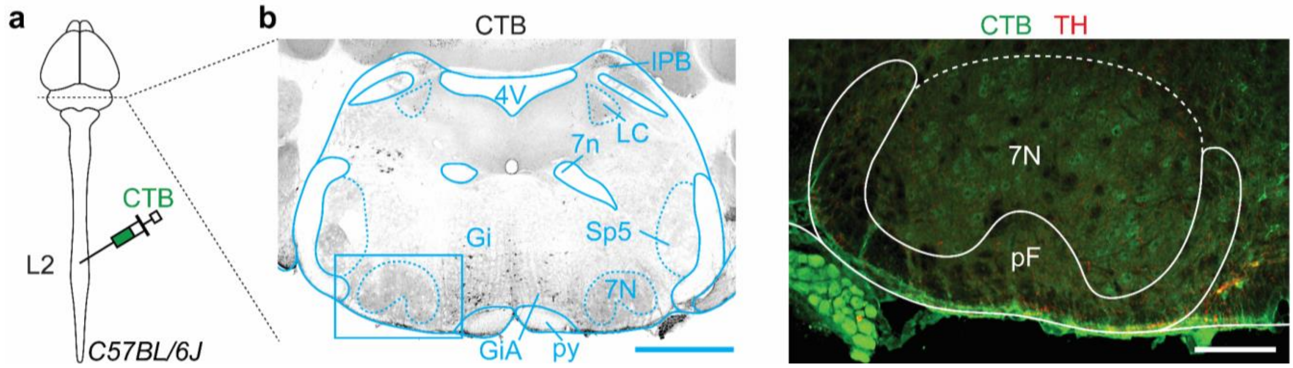


**Figure S6: Absence of spinally-projecting neurons in the pF. Related to Figure 4.**

**(a)** Experimental strategy for labelling spinally-projecting neurons: the retrograde tracer Cholera toxin B (CTB) is injected bilaterally at the second lumbar segment (L2) in a wild-type *C57BL/6J* adult mouse.

**(b)** Left: transverse section at the segmental level of the pF, showing CTB$^{+}$ spinally-projecting neurons in grey. Scale bar, 1 mm. Right: magnification of the boxed area showing that spinally-projecting neurons are not found in the pF but are located more medially and include catecholaminergic cell-types (expressing tyrosine hydroxylase, TH). Scale bar, 200 µm. Representative of n = 3 animals.

lPB: lateral para-brachial nucleus. LC: locus coeruleus; 7n: facial nerve. See Figure 1 for other abbreviations used.

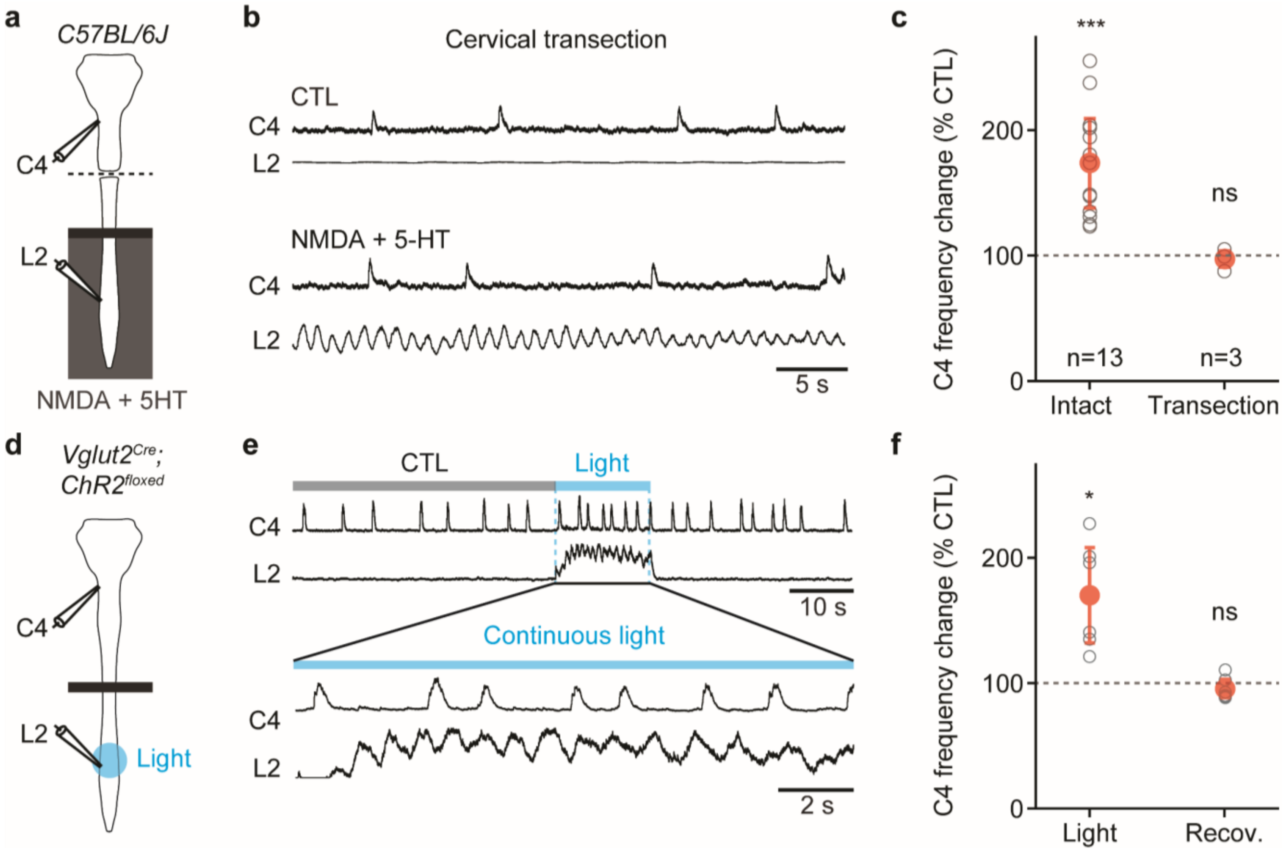


**Figure S7: Controls for *ex vivo* experiments and photoactivations of the glutamatergic lumbar spinal neurons. Related to Figure 5.**

**(a)** Schematics of the isolated *ex vivo* brainstem-spinal cord preparation from neonates (1-2 days). Respiratory- and locomotor-like activities are recorded using glass suction electrodes attached to the 4$^{th}$ cervical (C4) and the 2$^{nd}$ lumbar (L2) motor nerve roots respectively. The preparation is separated in two compartments using a Vaseline barrier (black bar). The lumbar spinal cord is superfused with control or locomotor drugs-enriched (NDMA and 5-HT, 10-14 µM each) artificial cerebrospinal fluid (aCSF) while the brainstem compartment remains in control aCSF. Here, and contrary to the experiments presented in Figure 5, preparations underwent a complete transection at the low cervical level.

**(b)** Recordings of respiratory- (C4) and locomotor-like (L2) activities of one representative animal before (CTL) and during perfusion of NMDA and 5-HT in the lumbar spinal cord compartment (only integrated traces are shown). The absence of change in C4 frequency following the transection rules out any leakage of NMDA and 5-HT from the spinal to the brainstem compartment.

**(c)** Quantification of the C4 frequency change during drug-induced locomotor-like activity as a percent change to the CTL condition, in both intact (n = 13) and sectioned (n = 3) preparations. Grey open circles are the means of individual preparations, and red circles are the means ± SD across n preparations.

**(d)** Similar preparations from *Vglut2$^{Cre}$; ChR2$^{floxed}$* neonates are used to photoactivate glutamatergic lumbar neurons with blue light delivered focally on the L2 segment.

**(e)** Recordings of C4 and L2 activities of one representative preparation before (CTL), during (Light) and after (OFF) a continuous 15 s light stimulation which triggers a substantial increase in respiratory frequency.

**(f)** Quantification of the C4 frequency change during photo-evoked locomotor-like activity expressed as a percent change to the CTL condition, in n = 6 preparations. In all graphs: ns, not significant. *, p<0.05; ***, p<0.001 (Wilcoxon matched-pairs tests).

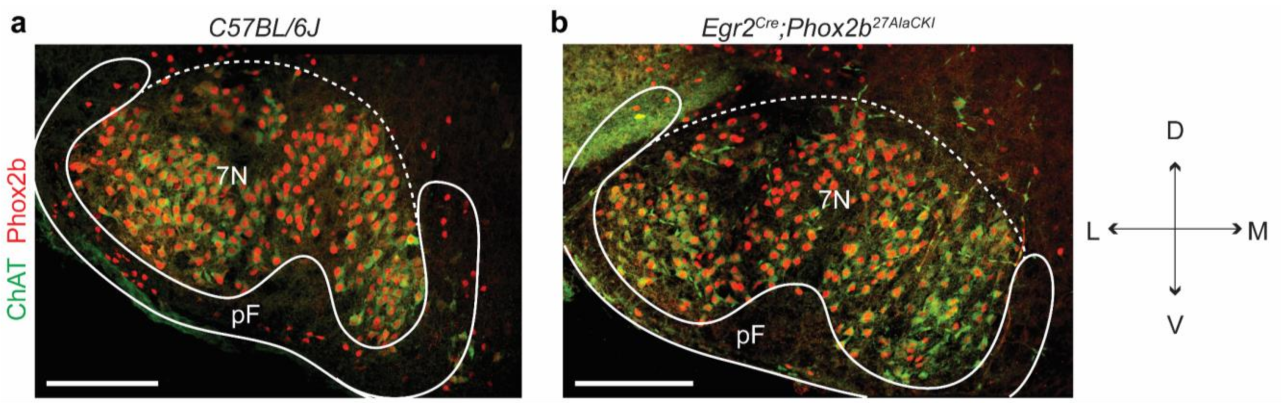


**Figure S8: Loss of RTN neurons in *Egr2$^{Cre}$;Phox2b$^{27AlaCKI}$* mutant neonates. Related to Figure 5.**

**(a)** Magnification of the pF area on a transverse brainstem section of a wild-type neonatal mouse. RTN neurons are detected as *Phox2b*-expressing interneurons located in the pF area lying ventrally, ventro-medially and ventro-laterally to the facial motoneurons (7N) which also co-express the Choline Acetyl Transferase (ChAT).

**(b)** Similar imaging from a *Egr2$^{Cre}$;Phox2b$^{27AlaCKI}$* neonate. Note the considerable loss of Phox2b-expressing RTN interneurons, as demonstrated previously in Ramanantsoa et al. 2011 and in Ruffault et al. 2015. Scale bars, 200 µm. Representative of 2 animals.

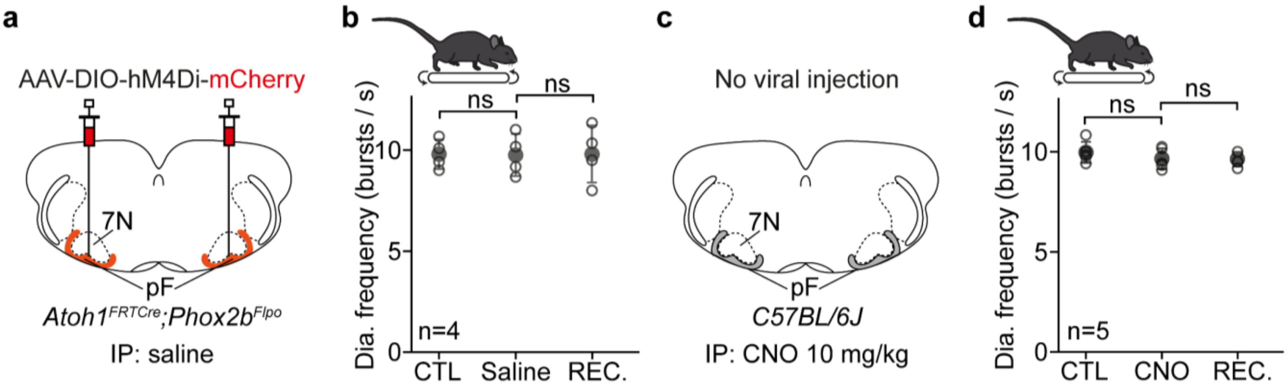


**Figure S9: Controls for chemogenetic experiments. Related to Figure 6.**

**(a)** Experimental strategy for the bilateral transfection of RTN$^{Phox2b/Atoh1}$ neurons with the inhibitory DREADD receptor hM4Di in an *Atoh1$^{FRTCre}$;Phox2b$^{Flpo}$* adult mouse. Injected mice were challenged to run on the motorized treadmill at 40 cm/s before, during and after intraperitoneal (IP) injection of saline solution.

**(b)** Quantifications of the diaphragm mean frequency before (CTL), during (Saline) and after (REC.) IP injection of saline. Grey open circles are the means of individual mice, and filled circles are the means ± SD across n mice. Saline injection alone does not alter the animals' capacity to increase its diaphragm frequency to normal values during treadmill running.

**(c)** Experimental strategy. Here, wild-type mice that did not receive any viral injection were challenged to run on the treadmill at 40 cm/s before, during and after IP injection of CNO at 10 mg/kg.

**(d)** Similar quantifications as above of the diaphragm mean frequency before (CTL), during (CNO) and after (REC.) IP injection of CNO. In all graphs: ns, not significant (Paired t-tests).

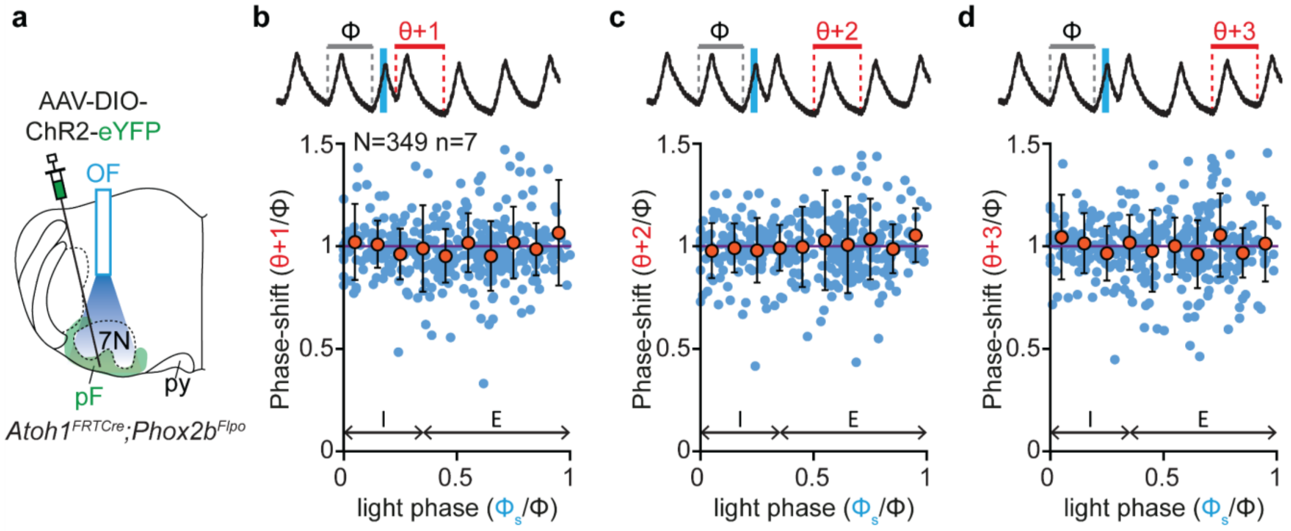


**Figure S10: Impact of photoactivating RTN$^{Phox2b/Atoh1}$ neurons on respiratory cycles following the light-perturbed cycle. Related to Figure 7.**

**(a)** Experimental strategy for photoactivating RTN$^{Phox2b/Atoh1}$ neurons unilaterally: an AAV-DIO-ChR2-eYFP is injected in the pF area of an *Atoh1$^{FRTCre}$;Phox2b$^{Flpo}$* adult mouse and an optic fiber (OF) is implanted above the injection site.

**(b)** Top: whole body plethysmography (WBP) recordings of respiratory cycles (inspirations are upwards, expirations are downwards) while a 50 ms pulse stimulation is applied. The cycle n+1 following the perturbed cycle is annotated on the trace (θ+1). Bottom: quantification of the perturbed cycle n+1 normalized to the control cycle (θ+1/ϕ) as a function of the light phase (light cycle normalized to the control cycle: $\phi_s/\phi$). Values < 1 (purple line) indicate a shortening of the perturbed cycle (see Figure 2 for details). Blue circles represent individual data from N = 349 random trials from n = 7 mice and orange circles are averages ± SD across all trials within 0.1 ms bins. Inspiration (I) and expiration (E) mean durations are annotated on the graph.

**(c, d)** Similar representations for cycles n+2 (θ+2, in c) and n+3 (θ+3, in d).
